# Structures of EHD2 filaments on curved membranes provide a model for caveolar neck stabilization

**DOI:** 10.1101/2025.06.05.658037

**Authors:** Elena Vázquez-Sarandeses, Vasilii Mikirtumov, Jeffrey K. Noel, Mikhail Kudryashev, Oliver Daumke

## Abstract

Caveolae are flask-shaped invaginations of the plasma membrane serving critical functions in mechano-protection and signal transduction. Caveolar dynamics, such as movement within the plasma membrane or endocytosis, relies on precise shaping of the highly curved caveolar necks. The dynamin-like EHD2 ATPase is proposed to form an oligomeric scaffold around the caveolar neck, but its detailed molecular action is poorly understood. Here, we employed cryo-electron tomography to elucidate structures of EHD2 filaments oligomerized on tubulated liposomes. EHD2 filaments form a highly curved membrane scaffold which stabilizes a sophisticated tubular membrane geometry with undulations along the tube’s axis. An amino-terminal sequence facilitates this geometry by inserting into the membrane, thereby acting as a spacer between adjacent filaments. Moreover, in endothelial cells lacking EHD2, caveolar necks become narrower and elongated. Our structural work provides the molecular framework for understanding EHD2 scaffold formation and its cellular function in caveolar dynamics.

## Introduction

The family of Eps15-homology domain-containing proteins (EHDs) are dynamin-related ATPases found exclusively in eukaryotes (Naslavsky & Caplan, 2011). The four mammalian EHD proteins have been associated with diverse cellular processes that require membrane remodeling and/or preservation of specific membrane shape (reviewed in (Bhattacharyya & Pucadyil, 2020)): EHD1 mediates endocytic recycling by regulating vesicle fission and membrane trafficking (Caplan *et al*, 2002; Deo *et al*, 2018; Pant *et al*, 2009; Sharma *et al*, 2010). Together with Rab11FIP5, rabenosyn-5, VPS45, and VIPAS39, EHD1 forms the FERARI complex which coordinates membrane fusion and fission in the Rab11-dependent recycling pathway (Solinger *et al*, 2020). EHD1 in cooperation with EHD3 also participates in ciliogenesis (Lu *et al*, 2015). Furthermore, EHD1 cooperates with the Bin-Amphiphysin-Rvs167 (BAR)-domain containing BIN1 protein during endocytic recycling (Pant *et al*., 2009) and during the formation of T-tubules (Demonbreun *et al*, 2015; Posey *et al*, 2014). Deletion of EHD3 in mice leads to enlarged hearts and abnormal cardiac function (Curran *et al*, 2014). In neurons, EHD4 was shown to localize to macropinosomes (Shao *et al*, 2002) mediating TrkA receptor uptake via macropinocytosis (Valdez *et al*, 2005). In non-neuronal cells, EHD4 regulates EHD1-mediated endosomal recruitment and vascular endothelial (VE)-cadherin membrane trafficking (Jones *et al*, 2020; Malinova *et al*, 2021).

EHD2 accumulates at injury sites in human myotubes and assists in the membrane repair process (Doherty *et al*, 2008; Marg *et al*, 2012). The best characterized function of EHD2, however, is linked to its localization at caveolae (Hoernke *et al*, 2017; Hubert *et al*, 2020; Ludwig *et al*, 2013; Matthaeus *et al*, 2020; Moren *et al*, 2019; Moren *et al*, 2012; Shah *et al*, 2014; Stoeber *et al*, 2012). Caveolae are small plasma membrane invaginations involved in mechano-protection and control of lipid homeostasis (Parton *et al*, 2020; Sotodosos-Alonso *et al*, 2023). EHD2 was proposed to stabilize caveolae by oligomerizing around their necks (Ludwig *et al*., 2013; Moren *et al*., 2012; Stoeber *et al*., 2012). Mutations in EHD2 leading to disturbed membrane binding, oligomerization, nucleotide binding or hydrolysis (Daumke *et al*, 2007) yield abnormal caveolar morphologies (Hoernke *et al*., 2017; Moren *et al*., 2012). In a mouse model deficient for EHD2, various cell types display detached caveolae and increased caveolar mobility (Hubert *et al*., 2020; Matthaeus *et al*., 2020), closely phenocopying the effect of EHD2 knockdown in cell culture (Moren *et al*., 2012). Accordingly, it was suggested that EHD2 restricts lateral diffusion and detachment of caveolae from the plasma membrane. In the EHD2 knockout mice, adipocytes were larger and contained more and expanded lipid droplets, concomitant with increased visceral fat deposits surrounding the organs (Matthaeus *et al*., 2020). Furthermore, in mesentery arteries, EHD2-dependent caveolae stabilization is required for mesentery relaxation via the endothelial nitric oxide synthase pathway (Matthaeus *et al*, 2019). Collectively, EHD2 appears to be a critical component for maintaining caveolae-associated signaling and uptake functions.

Biochemical and structural analyses provided insights into the mechanism of EHDs in membrane remodeling. The GTPase (G-) domain of EHDs surprisingly binds to adenine rather than guanine nucleotides (Daumke *et al*., 2007; Lee *et al*, 2005). When incubated with liposomes, EHDs form ATP-dependent oligomeric assemblies at the membrane surface, leading to liposome tubulation (Daumke *et al*., 2007; Deo *et al*., 2018; Melo *et al*, 2017; Melo *et al*, 2022; Pant *et al*., 2009; Shah *et al*., 2014). In the membrane-bound, oligomeric state, the slow ATPase activity of EHD proteins is moderately stimulated (Daumke *et al*., 2007; Deo *et al*., 2018; Melo *et al*., 2017). EHD1, but not EHD2 was shown to cleave preformed membrane tubules in an ATPase-dependent manner (Deo *et al*., 2018).

Crystallographic structural analyses indicated that the G-domain mediates stable dimerization via a unique interface in the dynamin superfamily (Supplementary Fig. 1). The adjacent helical domain is composed of sequences before and after the G-domain and contains the primary membrane-binding site at its tip (Daumke *et al*., 2007). The helical domain is connected via a long linker to the C-terminal EH domain (Fig. 1A), which mediates binding to linear peptide motifs containing an NPF (Asn-Pro-Phe) motif (Daumke *et al*., 2007; Kieken *et al*, 2009). In the crystal structure of the dimer, each EH domain is positioned on top of the opposing G-domain and binds to Gly-Pro-Phe (GPF) motif in the linker region between the helical and EH domain. The two helical domains protrude in parallel, representing the ‘closed’ conformation of EHDs (Supplementary Fig. 1A) (Daumke *et al*., 2007).

**Figure 1:**
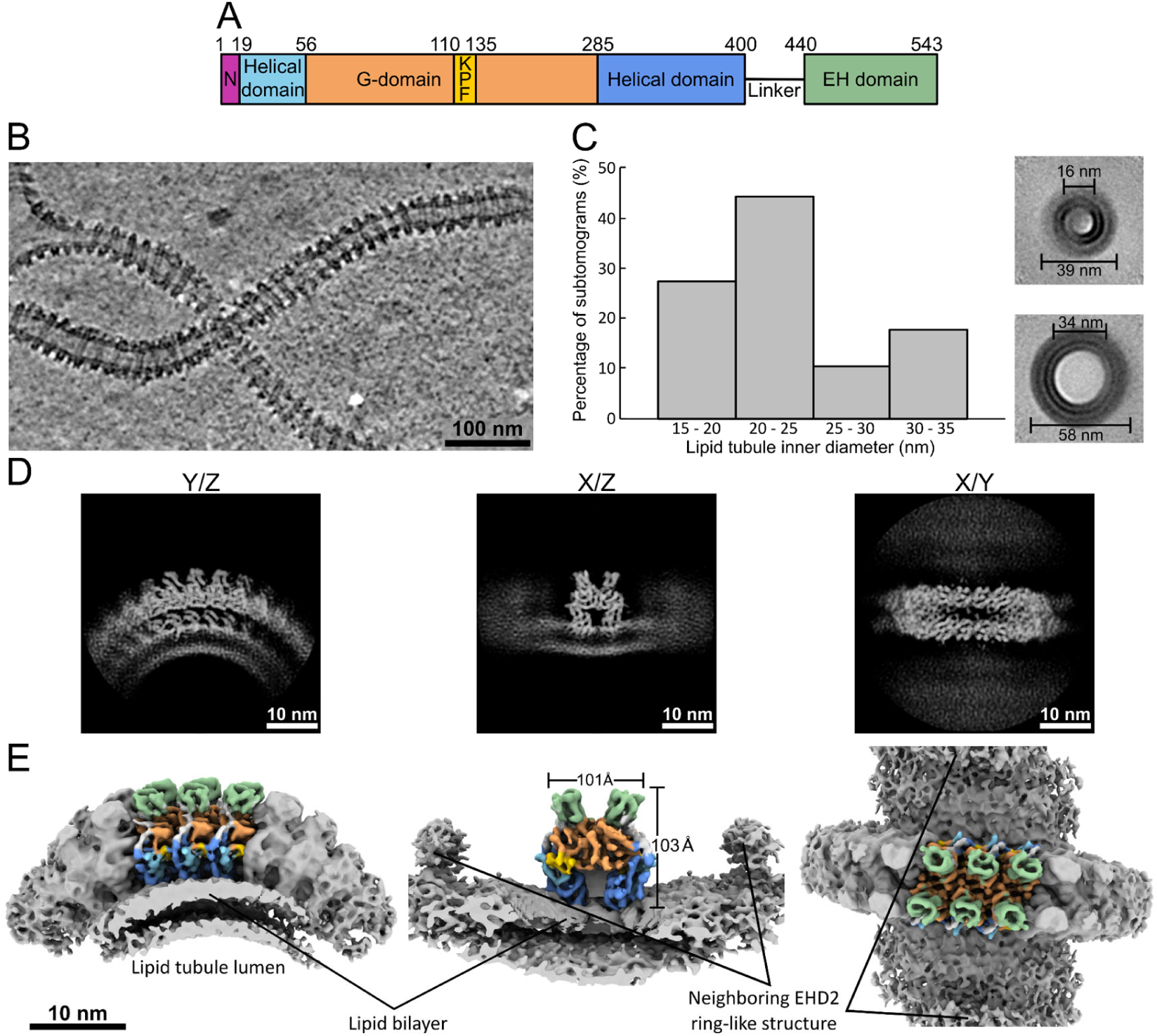
Structure determination of membrane-bound EHD2. **A)** Domain architecture of EHD2. Residue numbers refer to the mouse sequence. **B)** Central slice of a representative cryo electron tomogram of lipid tubules decorated with EHD2 ring-like oligomers. **C)** Distribution of particles according to the lipid tubules’ inner diameter. The lumen of the tubules, as measured in cross-sections of full 2D projections of subtomogram averages, is within a range of 16 – 34 nm. **D)** Projections of the resulting subtomogram average map of membrane-bound full-length EHD2. The view axis is indicated on top of each panel. **E)** Surface representation of the subtomogram average map. The view axis is the same as the panels above in D. The asymmetric unit, resolved at an average resolution of 6.7 Å, is colored according to **A**.

The crystal structure of an N-terminally truncated variant of EHD4 (EHD4^ΔN^) featured a large-scale 50° rotation of the helical domain relative to the G-domain compared to the EHD2 crystal structure. This arrangement was termed the ‘open’ conformation (Supplementary Fig. 1B) (Melo *et al*., 2017). While spectroscopic studies suggested that EHD2 recruitment to membranes occurs in the open state (Hoernke *et al*., 2017), a cryo-electron tomography (cryo-ET) derived subtomogram averaging structure (STA) indicated that membrane-bound EHD4^ΔN^ filaments feature a close conformation of the dimeric building block (Melo *et al*., 2022). Accordingly, it was proposed that EHDs exist as open dimers in solution and undergo a conformational rearrangement to the closed state when recruited to membranes and assembled into membrane-bound oligomeric filaments.

In the EHD4^ΔN^ filaments, oligomerization is mediated via three distinct interfaces: Interface-1 represents the EHD-specific dimerization interface also found in the open and closed dimeric crystal structures. Interface-2 involves a KPF (for Lys-Pro-Phe)-containing loop in the G-domain that assembles with the helical domain of the adjacent dimer (Melo *et al*., 2017; Melo *et al*., 2022). Finally, a conserved G-domain interface across the nucleotide-binding site (the G-interface) constitutes interface-3. Oligomeric EHD4^ΔN^ assembled into filaments of low curvature, namely a spontaneous curvature of ∼1/70 nm^-1^ (Melo *et al*., 2022). In contrast, EHD2 oligomerizes at membranes into ring-like structures of much higher curvature (Daumke *et al*., 2007). The molecular basis of the different assembly types in EHD4 and EHD2 is not known.

In the crystal structure of EHD2, a conserved N-terminal sequence stretch binds into a hydrophobic pocket at the periphery of the G-domain (Supplementary Fig. 1) (Shah *et al*., 2014). Electron paramagnetic resonance experiments indicated that the N-terminal stretch moves from the pocket in the G-domain into the membrane, representing a secondary membrane binding site (Shah *et al*., 2014). In turn, the KPF loop substitutes for the N-terminus in the hydrophobic G-domain pocket to generate interface-2 (Melo *et al*., 2017; Melo *et al*., 2022). The switch of the N-terminus was suggested to regulate EHD2 oligomerization at the caveolar neck (Hoernke *et al*., 2017; Shah *et al*., 2014). Also in EHD1, the deletion of the N-terminal residues caused defects in the stability of the membrane scaffold and affected its endocytic recycling activity (Deo *et al*., 2018). However, the structural role of the N-terminal sequence and its regulatory function have still remained unclear since the N-terminus was not fully resolved in the EHD2 crystal structures (Daumke *et al*., 2007; Shah *et al*., 2014) and was absent in the EHD4 construct used for crystal and STA-cryo-ET structure determination (Melo *et al*., 2017; Melo *et al*., 2022).

In the current study, we use cryo-ET and STA to determine the structure of membrane-bound full-length EHD2 reconstituted on highly curved lipid tubes. Compared to the previously described EHD4^ΔN^ filaments, we observe novel arrangements of EHD2 and EHD2^ΔN^ filaments on lipid tubes. The resolution of our STA structures allowed the fitting of available crystal structures including key oligomerization elements, providing novel insights into the assembly and membrane remodeling mechanism. The higher curvature of the EHD2 versus the EHD4^ΔN^ scaffold is achieved by a different assembly angle between the dimeric building blocks. By comparing the EHD2 and EHD2^ΔN^ filaments, we uncover a crucial role of the N-terminal sequence as a spacer, allowing the formation of a ring-like EHD2 scaffold stabilizing high membrane curvature at the caveolar neck.

## Results

### Structure determination of membrane-bound EHD2

We previously reported that mouse EHD2 binds to Folch liposomes and deforms them into tubules in an ATP-dependent fashion by forming oligomeric ring-like structures around them (Daumke *et al*., 2007). To structurally characterize the EHD2 assembly mode, we purified recombinantly-expressed full-length mouse EHD2 (Fig. 1A) and reconstituted it on membranes composed of Folch lipids. Using cryo-ET, we observed that EHD2 coated and occasionally tubulated liposomes in the absence of ATP, in line with our previous report (Shah *et al*., 2014). However, EHD2 failed to arrange into regular filaments in the absence of nucleotide (Supplementary Fig. 2A). In the presence of ATP, membrane tubulation was massively increased and ring-like EHD2 assemblies were found, confirming the ATP requirement for oligomerization (Fig. 1B and Supplementary Fig. 2B-E). In samples vitrified 2 h after their reconstitution with ATP (at this timepoint, about 90% ATP is hydrolyzed (Daumke *et al*., 2007)), oligomer coating was disrupted on many tubules, and the membrane tubules often displayed an irregular, thin appearance (Supplementary Fig. 3A). This is in line with our previous suggestions that ATP hydrolysis in EHD4^ΔN^ leads to oligomer disassembly (Melo *et al*., 2022) and highlights the scaffolding role of EHD2 in stabilizing membrane curvature.

We then collected 95 tilt-series of the ATP- and membrane-bound EHD2 filaments using a hybrid dose scheme with a high dose image at 0° tilt (Sanchez *et al*, 2020). In addition to the ring-like structures on highly-curved lipid tubules (Fig. 1B, Supplementary Fig. 2B and C), EHD2 also formed short oligomeric filaments on the surface of non-tubulated liposomes featuring low membrane curvature (Supplementary Fig. 2B, D). At transition areas, where lipid tubules emerged from the liposomes, short EHD2 filaments approached each other (Supplementary Fig. 2B, E), possibly in the process of oligomerizing into ring-like assemblies triggering membrane curvature.

To structurally characterize the membrane-bound EHD2 rings, we employed STA by defining subtomograms along the axis of the lipid tubules (Supplementary Fig. 4). Alignment and classification of the tubules revealed that their luminal diameters ranged from 16 to 34 nm (Fig. 1C and Supplementary Table 1). These radii are significantly smaller than those of the membrane tubes formed by EHD4^ΔN^, which were in the range of 34-87 nm (Melo *et al*., 2022). Several rounds of subtomogram classification and ‘subboxing’ (Castao-Dez *et al*, 2017) resulted in 75,439 lipid tube segments from an initial set of 14,491 subtomograms (Supplementary Fig. 4A). The final structure was refined by focusing on a segment within one ring, yielding an average resolution of 6.7 Å (Fig. 1D and E, Supplementary Fig. 4B-F). The asymmetric unit of the STA structure includes six monomers of EHD2 arranged into two dimers and two monomers corresponding to adjacent dimers within the filament (Fig. 1E, Supplementary Fig. 4F, Supplementary Table 2). Secondary structure elements could be clearly discerned in the map allowing an unambiguous identification of the G-, helical and EH domains: While the G-domains localize to the core of the filament, the helical domains extended towards the membrane and the EH domains locate on the top of the G-domains (Fig. 1E).

### Architecture of EHD2 filaments

A flexible fitting approach with the high-resolution EHD2 crystal structure was used to generate an atomic model of the EHD2 oligomer (Supplementary Fig. 5A). The G-domain covering residues 56-284 and the helical domains spanning residues 1-55 and 286-399 could be confidently modelled into the density (Fig. 2A). Clear density was evident for the KPF loop (residues 110-135) occupying a hydrophobic pocket in the G-domain (Fig. 2A), in which the N-terminus is buried in the crystal structure (Fig. 2B, Supplementary Fig. 1A). Similar to membrane-bound EHD4^ΔN^, oligomerized EHD2 was in the closed conformation (Fig. 2A, Supplementary Fig. 1A).

**Figure 2:**
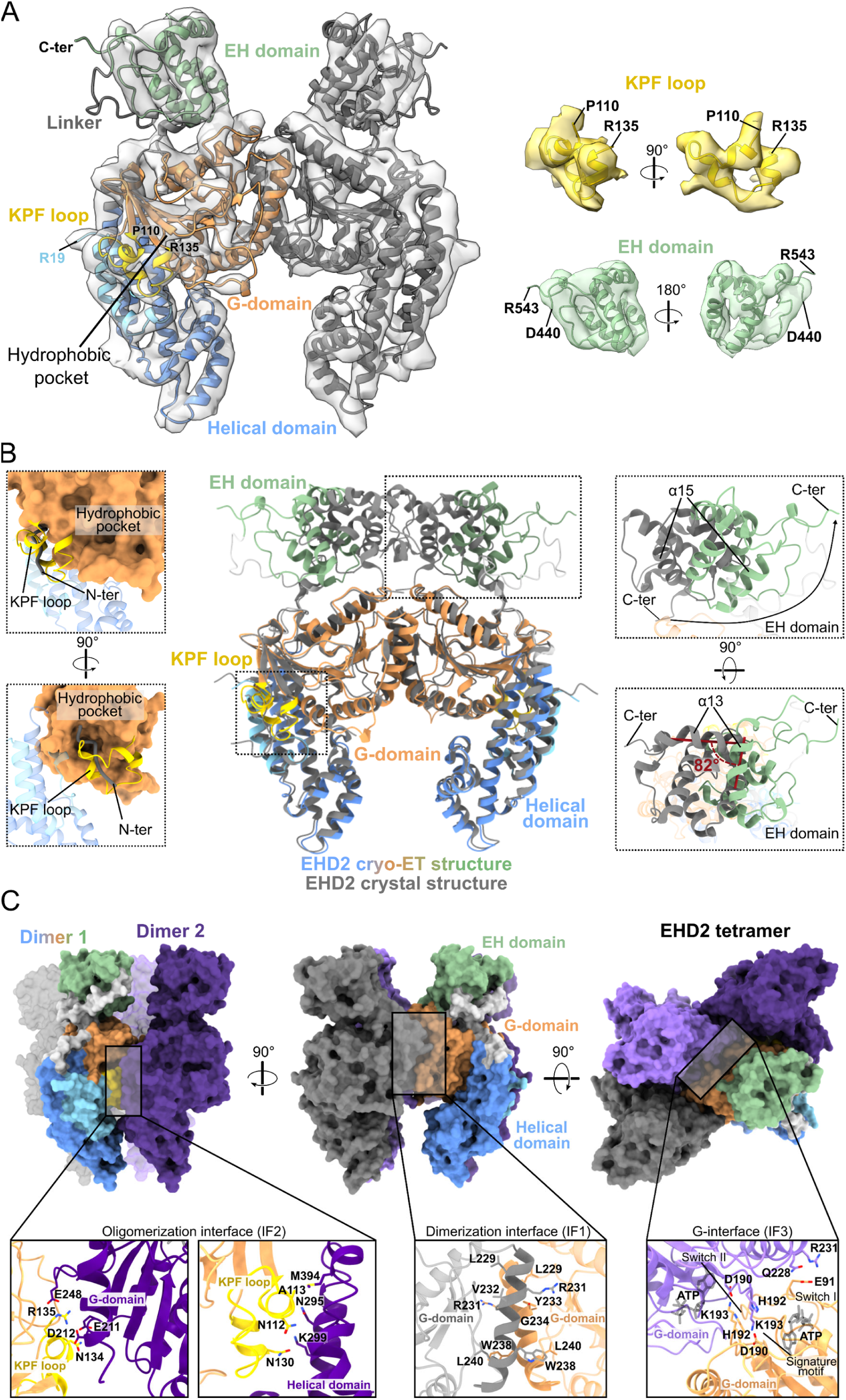
Structure of the EHD2 filaments. **A)** The resulting model of dimeric EHD2 fitted into the cryo-ET density. One monomer is colored according to the domains, and the other monomer is shown in gray. On the left, the fitting of the KPF loop and the EH domain in the density is highlighted. **B)** Superposition of the cryo-ET EHD2 structure (colored, only one dimer is shown) and the dimeric EHD2 crystal structure (gray, PDB: 4CID). Structural differences are highlighted and magnified in the dotted squares. The EH domain undergoes a large-scale rotation of ∼80°, which repositions the C-terminal tail to the outside of the oligomeric filament, thereby increasing the space between the EH domains in the EHD2 dimer. The KPF loop occupies the hydrophobic pocket of the G-domain (surface representation in the dotted square), where the N-terminus (gray) is buried in the crystal structure. **C)** Three interfaces drive oligomeric assembly and are highlighted in the central tetramer of the asymmetric unit. One dimer is formed by the monomer colored according to the domains and the monomer shown in gray. The other dimer is shown in two shades of purple. Front, side and top views correspond to the left, middle and right panels. Highlighted panels are magnified. The dimerization interface involves the G-domains (interface-1, IF1). The oligomerization interface (interface-2, IF2) is established between the KPF loop of one monomer and the helical and G-domains of the neighboring monomer from the adjacent dimer. The canonical G-interface (interface-3, IF3) is formed between the nucleotide binding pockets of opposing monomers from adjacent dimers.

In contrast to the G-domain and helical domain, the EH domains were more flexible and, consequently, the corresponding densities had a lower resolution of ∼7-8 Å (Supplementary Fig. 6). To obtain a reliable fitting, we calculated correlation scores of an EH domain model in 70 rotations equally distributed around a unit sphere with the excised EH domain map (Supplementary Fig. 6A). In the model with the highest correlation score, the C-terminal tail pointed towards the outside of the filament (Fig. 2A and B, Supplementary Fig. 6A, B). Contacts of EH domain helices α14 and α15 with the underlying G-domain stabilize the EH domain conformation in the oligomeric assembly (Supplementary Fig. 6C). Comparison of the EHD2 crystal structure and the STA-cryo-ET-based model reveals that the EH domains undergo a large-scale ∼80° rotation upon oligomerization, resulting in a shift towards the periphery of the filament (Fig. 2B). No density was apparent for the long, disordered linker between the EH domain and the helical domain, leading to an ambiguity in the connection of the domains. We assigned the EH domain to the helical domain situated directly below, since the connection of the domain termini was the closest in this way.

Three interfaces were previously defined in the EHD4^ΔN^ oligomer (Melo *et al*., 2022), which we also found in the EHD2 oligomer (Fig. 2C). The resolution of the EM map did not allow an accurate assignment of side chains, but with the EHD2 crystal structure as a basis, we included them in the model to highlight possible molecular interactions (Fig. 2C, Supplementary Fig. 4).

Interface-1 mediates EHD2 dimerization via the EHD-specific interface and mainly involves helices α6 from the G-domains of opposing monomers, which assemble in a symmetric fashion (Fig. 2C). Interface-2 and 3 drive the oligomerization of EHD2 dimers. Oligomerization interface-2 involves the KPF loop of one protomer and helices α8 and α12 from the helical domain of the adjacent dimer. Additional contacts not described for the EHD4^ΔN^ oligomer are formed between the KPF loop of one protomer and two short loops from the opposing G-domain (Fig. 2C). The canonical G-interface, designated interface-3, is built between residues surrounding the nucleotide binding pockets of opposing G-domains from adjacent dimers (Fig. 2C). The interface involves contacts of highly-conserved surface-exposed loops, such as switch I and II and the EHD-specific signature motifs. In analogy to other dynamin-related proteins (Daumke & Praefcke, 2016), this interface is likely responsible for the ATP-dependency of the assembly.

### The N-terminus acts as membrane-embedded spacer between filaments

When examining an unsharpened map, a distinctive density pattern was found directly next to the first ordered residue, Arg19, and reached towards the lipid bilayer while approaching the neighboring EHD2 filaments (Fig. 3A). In agreement with previous EPR data indicating membrane interaction of the N-terminus, we assigned this density to the N-terminal sequence and included it in our model. Notably, the N-terminal peptide is sufficiently long to cover the length of the density and insert with its N-terminal conserved hydrophobic and positively charged residues into the lipid bilayer (Fig. 3A-C).

**Figure 3:**
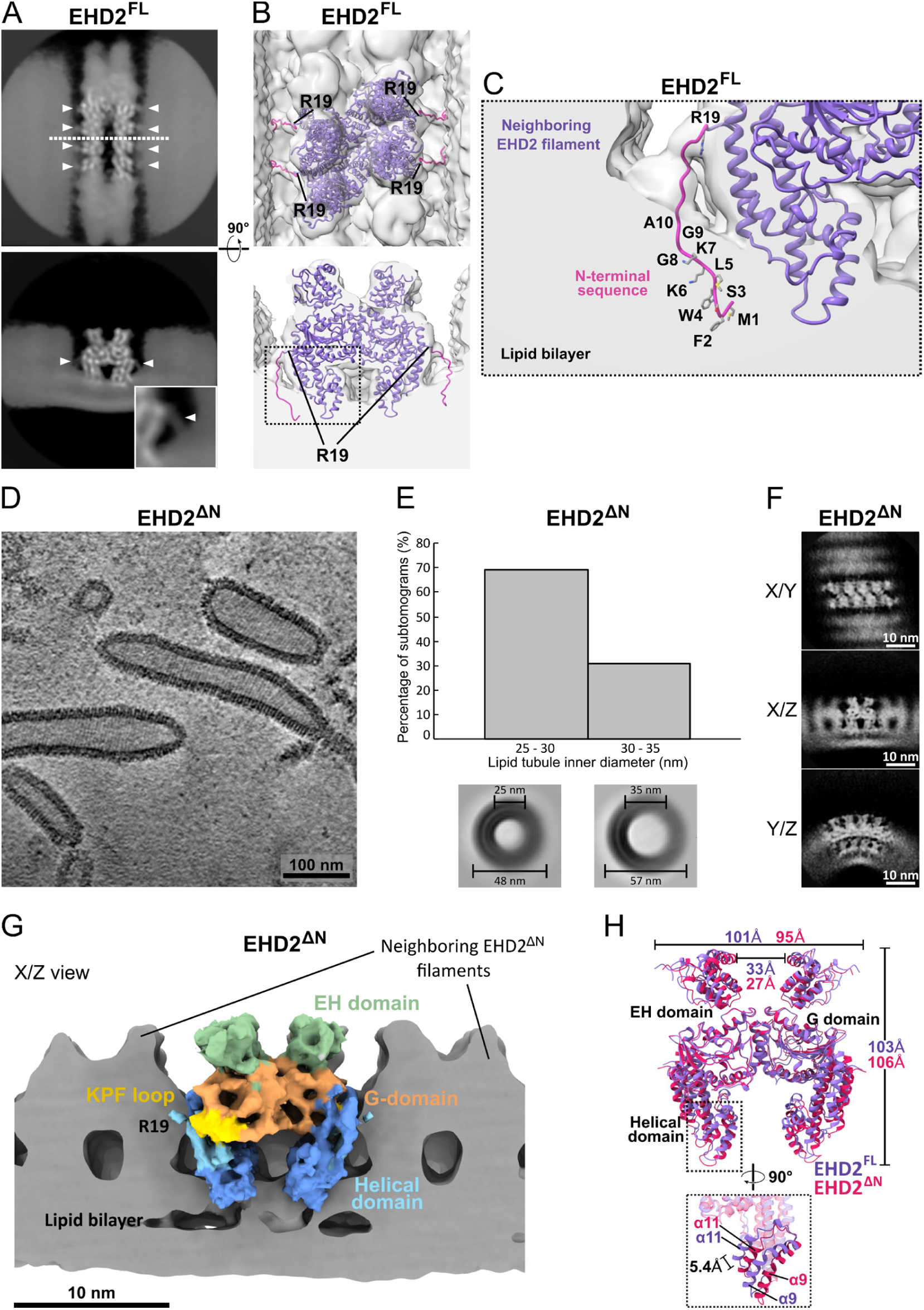
The N-terminus acts as a spacer between filaments. **A)** The unsharpened subtomogram averaging map reveals a pattern of low-resolution densities on the sides of the EHD2 filament (white arrowheads and magnified inlet). Top: Z view, bottom: Y view clipped at the white dashed line. **B)** 3D surface representation of the panels shown in A with the central EHD2 tetramer fitted in the density. The low-resolution densities emerge from the first modelled residue (Arg19) and reach towards the lipid bilayer. The N-terminal peptide (magenta) is long enough to cover the distance and insert into the lipid bilayer. **C)** Magnified view of the inlet highlighted in B. Hydrophobic and positively charged residues may insert into the outer leaflet of the lipid bilayer and come into proximity with the neighboring EHD2 filament. **D)** The absence of the N-terminus results in tightly packed oligomeric filaments around the lipid tubules. A central slice of a representative tomogram is shown. **E)** Distribution of particles according to lipid tubule inner diameter, measured in cross-sections of full 2D projections of subtomogram averages. The lipid tubules generated by EHD2^ΔN^ are more homogenous in terms of lumen diameter compared to full-length EHD2. **F)** Projections of the resulting subtomogram average map of membrane-bound full-length EHD2^ΔN^, obtained at an average resolution of 10.1 Å. Neighboring filaments in close proximity are observed in the Z and Y views. **G)** Surface representation of the subtomogram average map. The view axis (X/Z) is the same as the middle panel in F. The central tetramer is colored according to the domains. **H)** Superposition of full-length EHD2 (purple) and EHD2^ΔN^ (magenta) cryo-ET models. The deletion of the N-terminus results in a slightly higher structure with the EH domains moderately closer to each other. The switch-II region in the G-domain is shifted upwards by 4.5 Å and helices α9 and α11 from the helical domain are displaced by 5.4 Å (magnified). Note that the disordered linker between the helical and the EH domains is hidden for better visualization.

To further characterize the role of the N-terminus for the assembly, we reconstituted an N-terminally truncated EHD2 construct lacking the first 18 amino acids (residues 19-543, EHD2^ΔN^) on liposomes. We noticed that EHD2^ΔN^ filaments formed a more tightly packed coat on the surface of lipid tubes compared to full length EHD2 (Fig. 3D, compare to Fig. 1B). Accordingly, EHD2^ΔN^-coated lipid tubes were more homogenous in size, featuring tubular membrane diameters of 25-35 nm (Fig. 3E).

Employing the same experimental setup as for full-length EHD2, we collected 110 tilt series for EHD2^ΔN^-covered lipid tubes (Supplementary Table 2). We then determined the structure of EHD2^ΔN^ at an average resolution of 10.1 Å by STA using 17,204 particles subboxed from an initial set of 30,449 subtomograms (Supplementary Fig. 7). The asymmetric unit includes eight monomers of EHD2^ΔN^ which form four dimers (Fig. 3F, Supplementary Table 2, Supplementary Fig. 7). Similar to the EHD2 filaments, the EHD2^ΔN^ model was generated using flexible fitting and the crystal structure of dimeric EHD2 as a starting model (Supplementary Fig. 5B).

As expected, the low-resolution density corresponding to the N-terminal residues in the EHD2 filaments was absent in the EHD2^ΔN^ map (Fig. 3G). The conformations of EHD2 and EHD^ΔN^ appeared overall similar in the filaments. Minor structural rearrangements of α9 and α11 in the helical domains were observed and the EH domains moved 6 Å closer to each other in EHD^ΔN^ compared to EHD2 (Fig. 3H). Also, the assembly mode of the EHD2^ΔN^ filaments via the three oligomerization interfaces was not perturbed.

Strikingly, EHD2^ΔN^ filaments assembled into tightly packed filaments rather than ring-like structures (Fig. 3D, compare to Fig. 1B). Similar to the EHD4^ΔN^ filaments, peripheral helices α1a, α1b and α2 in the G-domain contacted each other across adjacent filaments (Fig. 3F, G), whereas they were 70 Å apart in the EHD2 full-length assemblies (Fig. 4A, right). These observations indicate a function of the membrane-inserted N-terminus as a spacer between adjacent filaments required for the formation of distinct ring-like structures.

**Figure 4:**
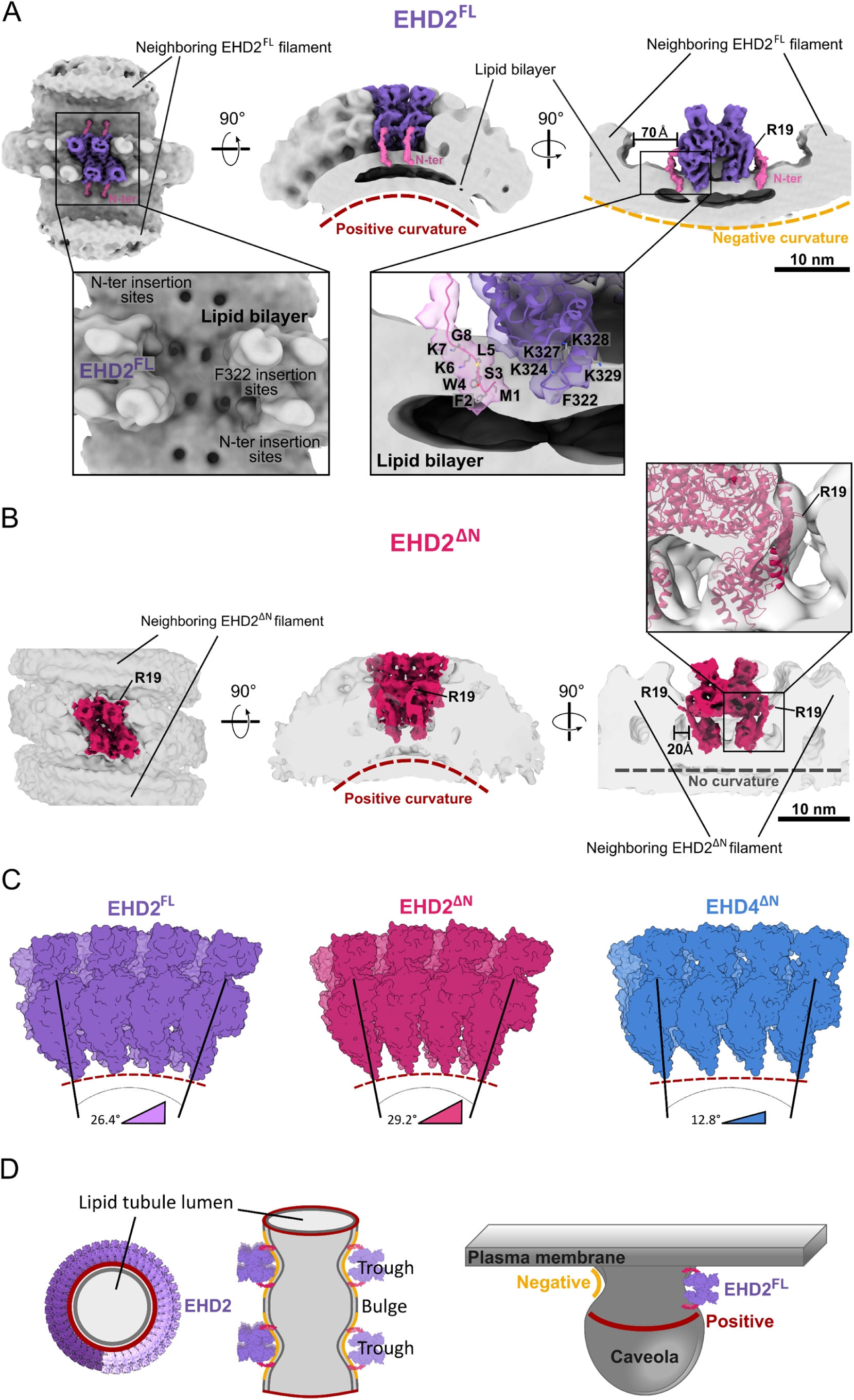
EHD2 filaments stabilize a membrane geometry reminiscent of the caveolar neck. **A)** 3D surface representation of the subtomogram average map including the lipid bilayer. The central tetramer of the asymmetric unit is highlighted in purple. The low-resolution density of the map where the N-terminus has been modeled is shown pink. Left: top view of an EHD2 ring-like structure. Middle: side view showing positive curvature of the membrane tubule stabilized by the EHD2 filament. Right: front view showing negative membrane curvature in undulations along the tubule’s axis. Magnified inlets show how EHD2 inserts the tip of the helical domain and the first residues of the N-terminus in the outer leaflet of the lipid bilayer. Left: The central tetramer is hidden to show how the helical domain and the N-terminus penetrate the membrane. Right: The residues involved in membrane binding are indicated in one monomer. **B)** 3D surface representation of the EHD2^ΔN^ subtomogram average map including the lipid bilayer. The central tetramer of the asymmetric unit is highlighted in magenta. Left: top view showing oligomeric filaments in close vicinity. Middle: side view showing positive membrane curvature. Right: the deletion of the N-terminus renders the lipid bilayer flat. The magnified inlet shows the absence of extra densities emerging from Arg19, which points toward the neighboring filament. **C)** EHD2 and EHD2^ΔN^ filaments (purple and magenta, respectively) are more curved than EHD4^ΔN^ filaments (blue, PDB: 7SOX). The dashed red lines indicate the positive curvature of the lipid bilayer. The side view of an octamer is shown for each protein. **D)** Schematic representation of the *in vitro* membrane tubulation activity of EHD2 and its relation to the suggested localization at the neck of caveolae, where positive and negative membrane curvature co-exist in a similar fashion.

### EHD2 filaments stabilize a tubular membrane geometry with undulations

Clear density for the lipid bilayer allowed us to characterize the membrane-binding mode of EHD2 (Fig. 4). EHD2 inserts Phe322 at the tip of the helical domain into the outer leaflet of the lipid bilayer which would be expected to induce membrane curvature by increasing the surface area of the outer membrane leaflet (Fig. 4A). In addition, positively charged residues, such as Lys324, and Lys327-329, which were previously shown to contribute to membrane binding, are in close contact to the bilayer (Fig. 4A) and likely mediate the phosphatidyl-inositol(4,5)bisphosphate specificity of EHD2 (Daumke *et al*., 2007; Melo *et al*., 2017). Moreover, the proposed position of the N-terminal stretch close to the membrane (Fig. 3A-C, Fig. 4A) supports its role as a secondary membrane-binding site (Shah *et al*., 2014) (Fig. 4A).

In the EHD2 full-length filaments, the membrane surface along the tubule’s axis showed undulations, with the EHD2 filament sitting in the troughs of the undulations (Fig. 4A, D). In this way, positive and negative membrane curvature is generated along the tubule’s axis (Fig. 4A, D). In contrast, no undulations were observed for EHD2^ΔN^ filaments, resulting in a flat lipid bilayer along the tube axis (Fig. 4B). Notably, the membrane geometry stabilized by EHD2 filaments resembles the membrane architecture of the caveolar neck, where also a combination of positive and negative membrane curvature is found (Fig. 4D).

The angle between two assembling EHD2 dimers was ∼26° (Fig. 4C). The N-terminal deletion did not grossly alter the assembly angle between two adjacent dimers (∼26° for full-length EHD2 versus ∼29° for EHD2^ΔN^) resulting in a similar diameter of the underlying membrane tube (Fig. 4C). In stark contrast, the assembly angle was only ∼13° in the EHD4^ΔN^ filaments (Melo *et al*., 2022). The higher assembly angle leads to the newly observed G-domain-G-domain contacts involving the KPF-loop (Fig. 2C) and underlies the smaller diameter of EHD2 rings which, accordingly, stabilize higher membrane curvature compared to EHD4^ΔN^ filaments.

### The role of EHD2 in stabilizing membrane curvature at the neck of caveolae

To relate the structural analysis of the reconstituted EHD2 to its cellular role at caveolae, we examined caveolar morphology in Human Umbilical Vein Endothelial cells (HUVEC) in the presence and absence of EHD2, using small interfering (si)RNA-mediated knockdown. Electron tomograms of semi-thin sections from resin-embedded HUVECs (obtained from (Matthaeus *et al*., 2019)) were recorded at room temperature, and both the length and the width of the caveolar bulbs and the necks were examined (Fig. 5A and Supplementary Table 3).

**Figure 5:**
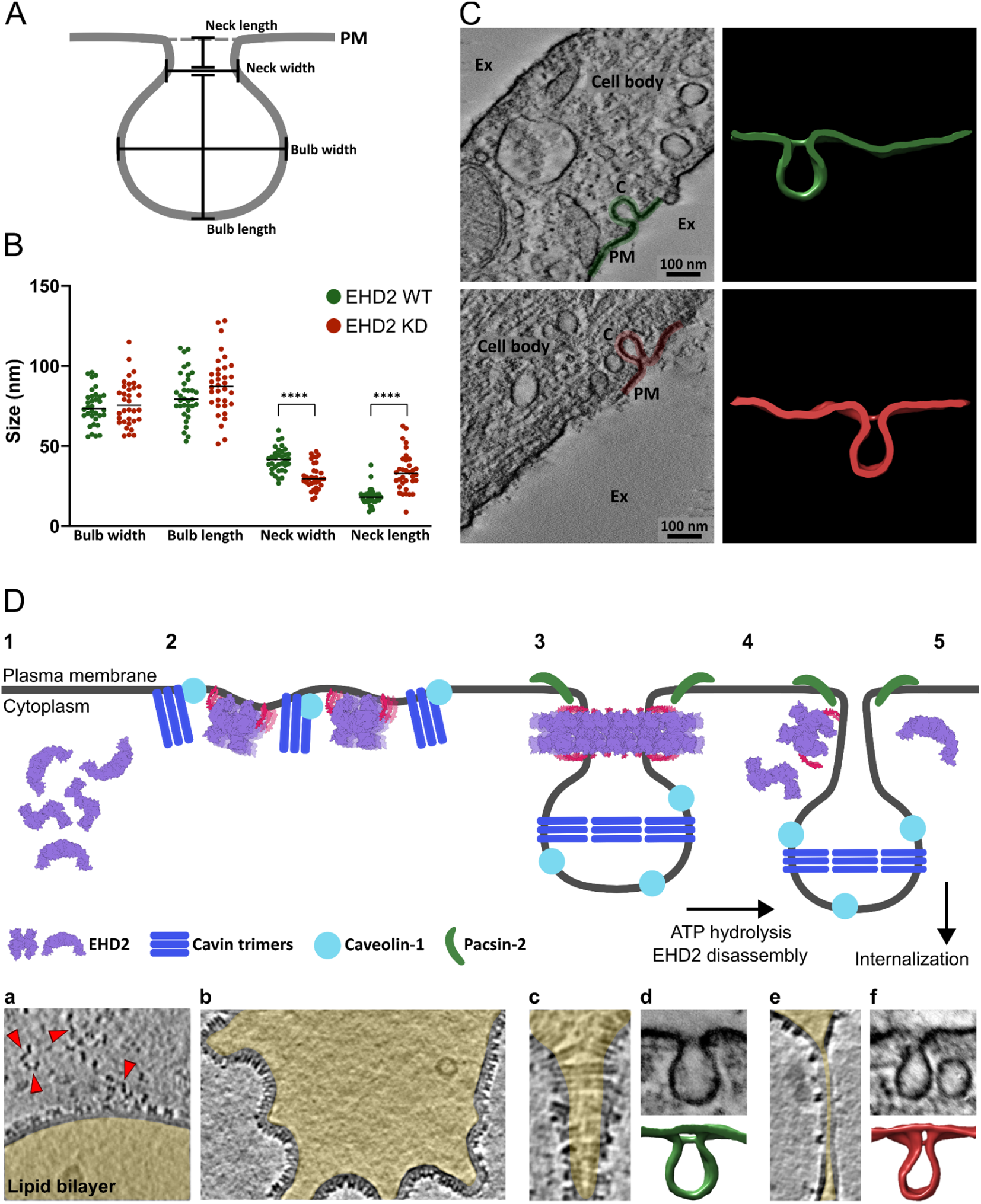
EHD2 serves as a scaffold to direct proper caveolae neck morphology. **A)** Schematic representation caveolae, indicating how their morphology was evaluated. **B)** Analysis of caveolae morphology in the presence (green) and absence (red) of EHD2. The deletion of EHD2 results in narrower and vertically elongated caveolar necks. The bulbs of caveolae remain unaffected. p ≤ 0.0001****. **C)** Central slices of representative room-temperature tomograms of EHD2 WT and EHD2 knockdown HUVEC cells. The 3D surface of a segmented average caveola is shown; PM: plasma membrane, Ex: extracellular space. **D)** Proposed model for EHD2 function at caveolae (top). 1) EHD2 exists as an open dimer in solution. 2) EHD2 is recruited to flat caveolae at the plasma membrane (Matthaeus *et al*., 2022) and undergoes a conformational change towards the closed oligomeric conformation. The N-terminus acts as secondary membrane binding site. Cavins and caveolins also found at flat caveolae initiate budding (Matthaeus *et al*., 2022), which might be further assisted by EHD2. 3) At the highly curved neck of caveolae, EHD2 forms ring-like oligomeric scaffolds. By generating and stabilizing both positive and negative curvature, EHD2 oligomers shape the caveolar neck, possibly assisted by the F-BAR domain protein PACSIN2 and other EHD2-binding partners like EHBP1 (Matthaeus *et al*., 2022). 4) ATP hydrolysis driven EHD2 disassembly and detachment from the plasma membrane destabilizes the membrane neck, leading to thinning of the neck. 5) Loss of the EHD2 scaffold allows internalization of caveolae. At the bottom, electron micrographs of reconstituted EHD2 samples or caveolar morphologies from HUVECs with related membrane geometries to the individual steps of the model are displayed; the vesicle’s/tube’s lumen is colored in yellow. a - ATP-bound EHD2 oligomers on a flat membrane surface; adjacent non-membrane-bound EHD2 dimers appear to be in an extended open conformation (red arrowheads, see Suppl. Fig. 1-3); b - Flat membrane surface decorated with ATP-bound EHD2 oligomers (see Suppl Fig. 2); c - EHD2 rings on a tubulated membrane tube of varying diameter (see Suppl. Fig. 2); d - Caveola with a regular membrane neck (see Fig. 5c); e – Thinned membrane tube in a reconstituted EHD2 samples after ATP hydrolysis (see Suppl. Fig. 3); f – Caveola featuring a thin neck in the absence of EHD2 (see Fig. 5c).

In agreement with previous data (Matthaeus *et al*., 2019), knockdown of EHD2 resulted in 2-fold increased caveolae detachment from the plasma membrane (Supplementary Table 3). The absence of EHD2 did not affect the shape of caveolar bulbs. However, the necks of caveolae in EHD2 knockdown cells were significantly narrower and vertically elongated (Fig. 5B, C): In HUVECs, the caveolar necks were 41 ± 8 nm wide and 18 ± 5 nm long, whereas they were 30 ± 8 nm wide and 33 ± 9 nm long in EHD2 knockdown HUVECs (Fig. 5C). These results are consistent with a model in which EHD2 filaments stabilize a defined diameter of the caveolar necks, therefore preventing their detachment from the plasma membrane.

## Discussion

Here, we elucidate the structural basis of EHD2 scaffold assembly and show how its assembly stabilizes highly-curved membrane tubes mimicking the caveolar neck. We uncover a role for the N-terminal sequence as a spacer between EHD2 filaments, preventing their aggregation and allowing the formation of ring-like structures. Our structural study sheds light on EHD2’s mechanisms as a stabilizer of the caveolar neck.

Membrane-bound oligomeric filaments, such as those of dynamin (Kong *et al*, 2018), Drp1 (Peng *et al*, 2025) or OPA1 (Nyenhuis *et al*, 2023; von der Malsburg *et al*, 2023), frequently feature helical symmetry. This enables straightforward “particle picking” in 2D and refinement with helical symmetry, often leading to high resolution reconstructions. In contrast, membrane-bound EHD2 samples were not helical and more heterogeneous, requiring application of cryo-ET and STA analyses to determine their structures. At an average resolution of ∼6.7 Å, the predominant α-helices in EHD2 can be resolved, while amino acid side chains are not visible. For the more flexible and therefore less well-defined regions, such as the EH-domain and especially the N-terminal sequence stretch, the domain orientation or the course towards the membrane could be deduced. By fitting the available EHD2 crystal structure into the map, we obtained a quasi-atomic model of the EHD2 filaments on membrane tubules allowing us to deduce molecular insights into the assembly mechanism.

EHD2 oligomerization is driven by the formation of the interfaces previously described for the EHD4 filaments: the EHD-specific dimerization interface in the G-domain, the oligomerization interface formed between the KPF-loop from the G-domain in one dimer and the helical domain of the neighboring dimer and the canonical G-interface, which involves the surface across the nucleotide binding pockets of neighboring dimers. In the EHD2 crystal structures, the C-terminus of the EH domain folds back to the G-domain blocking the formation of the G-interface and oligomerization in an auto-inhibitory fashion (Daumke *et al*., 2007; Shah *et al*., 2014). In contrast, in the EHD2 filament structure, the EH domains undergo a large-scale movement which repositions the C-terminal tail towards the outside, thereby relieving auto-inhibition of the G-interface. The NPF binding pockets of the EH domains point towards the inside of the filament and may only be partly accessible for peripheral interactions with described NPF-domain containing binding partners, such as EHBP1, PACSIN2, or MICAL-L1 (Giridharan *et al*, 2013; Guilherme *et al*, 2004; Senju *et al*, 2011). Thus, during complex formation, the EH domain may need further reorientation to direct the NPF-interaction site to their binding partners.

EHD2 dimers were found to oligomerize in the closed conformation on the lipid surface, with the tip of the helical domains inserting into the membrane. On membranes of low curvature, short oligomeric filaments are heterogeneously oriented, likely since their curvature does not match the curvature of the underlying membrane. On highly curved lipid tubules, EHD2 filaments adopt regular ring-like shapes, suggesting that high membrane curvature drives the formation of regular EHD2 filaments. In turn, EHD2 generates membrane curvature by inserting Phe322 at the tip of the helical domain into the outer leaflet of the membrane bilayer, as demonstrated by mutagenesis and EPR experiments (Daumke *et al*., 2007; Shah *et al*., 2014). Furthermore, the EHD2 filament acts as a curved membrane scaffold superimposing its curvature on the underlying membrane. Thus, EHD2 scaffold formation and membrane curvature generation are intimately intertwined. Compared to the previously described EHD4^ΔN^ filaments, the angle between assembling dimers is larger in EHD2 filaments, resulting in a higher curvature of the EHD2 scaffolds. This difference angle does not depend on the N-terminus of EHD2, as the EHD2^ΔN^ scaffold stabilizes a similar curvature compared to EHD2. Instead, the difference seems to reflect an intrinsic oligomerization property of individual EHD homologues, for example by unique interactions of the KPF-loop in the oligomer. The different filament curvatures may be adapted to the specific cellular sites of action: While EHD4 acts on larger macropinosomes of low membrane curvature, the EHD2 scaffold at the caveolar neck must stabilize a higher membrane curvature.

In addition to the curvature of the membrane tubule, EHD2 oligomers also induce undulations of the membrane surface along the tubule’s axis. Compared to previous models for EHD1 and EHD2 (Campelo *et al*, 2010; Daumke *et al*., 2007; Deo *et al*., 2018), the membrane-bound EHD2 filaments rest on the troughs, not on the ridges of the undulations. This specific membrane architecture depends on the N-terminus of EHDs. In the auto-inhibited EHD2 structure, the N-terminus of EHD2 is present in a hydrophobic pocket of the G-domain. In the EHD2 filament, we assigned it to a density reaching along the helical domain towards the membrane. This assignment is supported by our previous EPR experiments indicating membrane insertion of the N-terminus (Shah *et al*., 2014). Furthermore, the low-resolution density at the periphery of the G-domain was absent when the N-terminal stretch was deleted. The N-terminal truncation had a drastic effect on the overall arrangement of EHD2 filaments. Instead of the more or less regularly spaced rings, filaments now tightly approached each other, similar to the filaments described for EHD4^ΔN^ (Melo *et al*., 2022). Deletion of the EHD2 N-terminus also prevented membrane bulging along the axis of the membrane tubule. Also, the increased membrane recruitment of a transiently overexpressed EHD2^ΔN^ mutant (Hoernke *et al*., 2017; Shah *et al*., 2014) can be explained by a tighter assembly geometry. A related role of the N-terminus as an architectural element was shown for EHD1, as its deletion led to defects in scaffolding, scission and endocytic recycling (Deo *et al*., 2018). We therefore propose that upon membrane recruitment, the conserved N-terminal sequence switches from the G-domain into the membrane to act as a molecular spacer posing steric constraints between EHD oligomers required for their proper assembly and function.

In the cellular environment, the N-terminal spacer may prevent the aggregation of neighboring EHD2 helices and instead favor the formation of a single EHD2 ring at the caveolar neck. The caveolar neck has a related geometry to the EHD2-coated membrane undulations observed in our cryo-ET reconstruction, featuring a constriction with positive and negative membrane curvature (Kozlov & Taraska, 2023; Ludwig *et al*., 2013; Parton *et al*., 2020). Furthermore, the diameter range of the lipid tubules in our *in vitro* reconstitution is similar to the diameter of the caveolar neck (Hubert *et al*., 2020; Matthaeus *et al*, 2022; Parton *et al*., 2020; Sotodosos-Alonso *et al*., 2023). In an endothelial cell line, we observed significant alterations in caveolar morphology upon EHD2 knockdown only for the caveolar necks, not for the caveolar bulbs. The few caveolae confined to the plasma membrane in EHD2 knockout cells exhibited narrower and more elongated necks compared to those of wild-type cells. This observation corroborates our idea that the reduced overall caveolar mobility (Hubert *et al*., 2020; Matthaeus *et al*., 2020; Moren *et al*., 2012; Stoeber *et al*., 2012) is related to alterations at the caveolar neck and that EHD2 serves as a membrane-stabilizing scaffold at the caveolar neck. Moreover, non-invaginated caveolae (Matthaeus *et al*., 2022) may contain short oligomeric EHD2 filaments, and their oligomerization may support the formation of mature caveolae.

By integrating our structural results on EHD2 with previous structural and biochemical data, we suggest a refined model of the EHD2 cycle during caveolar function (Figure 5D). In this model, ATP-bound EHD2 dimers are recruited in the open conformation (Hoernke *et al*., 2017) to flat caveolae at the plasma membrane and assemble into short oligomeric structures. When caveolae invaginate, the short EHD2 oligomers may transition to the closed conformation and form rings surrounding the caveolar neck. Upon membrane binding, the N-terminus switches from the G-domain into the membrane and acts as a spacer preventing the uncontrolled oligomerization of EHD2 filaments. In this way, a single EHD2 ring stabilizes the caveolar neck at its thinnest position to a defined diameter. Oligomerization-dependent ATP hydrolysis may set an intrinsic timer for destabilizing the G-interface, prompting the disassembly of the EHD2 oligomer (Deo *et al*., 2018; Hoernke *et al*., 2017; Melo *et al*., 2022). In the absence of an EHD2 scaffold, the caveolar neck may become unstable, promoting the detachment of caveolae from the plasma membrane. In the cytosol, the EHD2 dimer may switch back to the open conformation to resume a new reaction cycle.

In summary, we used cryo-ET and STA to characterize the structural basis of EHD2 filament assembly and the role of these filaments in stabilizing highly curved membrane tubes mimicking the caveolar neck. Our study lays the groundwork for future *in situ* approaches aimed at resolving EHD2 structures at native caveolae, potentially capturing additional aspects of the cellular context, such as the presence of interaction partners or the influence of certain lipids enriched at caveolae.

## Materials and Methods

### Protein purification

Mouse EHD2 full-length (residues 1-543) and EHD2^ΔN^ (residues 19-543) constructs were expressed in *E. coli* (BL21(DE3)-Rosetta2 strain) from a modified pET28 vector as N-terminal His6-tag fusions followed by a PreScission protease cleavage site (according to (Daumke *et al*., 2007). Expression plasmids were transformed in *E. coli* host strain BL21(DE3)-Rosetta2 (Novagen). Cells were grown shaking at 37 °C in TB medium. Protein expression was induced by the addition of 40 μM isopropyl-β-D-thiogalactopyranoside (IPTG) at an optical density of 0.6, followed by overnight incubation with shaking at 18 °C. Cells were harvested by centrifugation (4,500 rounds per min (rpm), 20 min, 4 °C) and pellets were resuspended in resuspension buffer (50 mM HEPES/NaOH pH 7.5, 400 mM NaCl, 25 mM imidazole, 2.5 mM β-mercaptoethanol, 250 µM Pefabloc, 1 µM DNase I). Lysis was carried out using a microfluidizer. After centrifugation (20,000 rpm, 40 min, 4 °C), cleared lysates corresponding to the soluble protein fraction were applied to a Ni-NTA column. The column was extensively washed using washing buffer I (20 mM HEPES/NaOH pH 7.5, 700 mM NaCl, 30 mM imidazole, 2.5 mM β-mercaptoethanol, 1 mM ATP, 10 mM KCl) and washing buffer II (20 mM HEPES/NaOH pH 7.5, 300 mM NaCl, 25 mM imidazole, 2.5 mM β-mercaptoethanol). The protein was eluted using elution buffer I (20 mM HEPES/NaOH pH 7.5, 300 mM NaCl, 300 mM imidazole, 2.5 mM β-mercaptoethanol). For His-tag cleavage, 150 μg of PreScission protease were used per 5mg of EHD2 construct. The protein sample was dialyzed overnight at 4 °C against dialysis buffer (20 mM HEPES/NaOH pH 7.5, 300 mM NaCl, 2.5 mM β-mercaptoethanol) for imidazole removal and then re-applied to the Ni-NTA column for His-tag removal. The protein was eluted in two steps of increasing imidazole concentration using washing buffer II and elution buffer II (20 mM HEPES/NaOH pH 7.5, 300 mM NaCl, 50 mM imidazole, 2.5 mM β-mercaptoethanol). Concentrated protein was injected into a Superdex200 gel filtration column, previously equilibrated with SEC Buffer (20 mM HEPES/NaOH pH 7.5, 300 mM NaCl, 2.5 mM β-mercaptoethanol, 2.5 mM MgCl_2_). A second run of size exclusion chromatography was performed as a polishing step. Fractions containing EHD2 constructs were pooled, concentrated and flash-frozen in liquid nitrogen.

### Liposome preparation

Folch fraction I bovine brain lipids (Sigma) were dissolved in chloroform at a concentration of 25 mg/ml. To form the liposomes, 50 μl of the lipid solution were mixed with 200 μl of a Chloroform/Methanol (3:1 v/v) mixture and dried under an argon stream and inside a desiccator. The lipids were resuspended in liposome buffer (20 mM HEPES/NaOH pH 7.5, 300 mM NaCl, 1 mM β-mercaptoethanol) and sonicated in a water bath for 30 seconds.

### Cryo-electron tomography

For the generation of protein-decorated lipid tubules, 80 μM of the indicated EHD2 construct diluted in tubulation buffer (20 mM HEPES/NaOH pH 7.5, 300 mM NaCl, 0.5 mM MgCl_2_) was incubated with 1.125 mM ATP for 5 min at room temperature. Afterwards, Folch liposomes diluted in liposome buffer were added to yield a final concentration of 2 mg/ml. The sample was further incubated for 10 min at room temperature and, prior to plunge-freezing in liquid ethane, 5 nm colloidal gold was added at a 1:40 ratio (v/v). For apo conditions, the 5 min incubation with ATP was omitted. Glow-discharged carbon-coated copper Quantifoil 2/2 grids were spotted with 4 μl of sample, back-blotted for 3 seconds and plunge-frozen using a Vitrobot Mark II device. Tilt series were acquired using a TFS Titan Krios G3 electron microscope equipped with a Gatan K3 detector and a Bioquantum energy filter and operated at 300 kV in zero-loss mode. The tilt series were collected using the software SerialEM (Mastronarde, 2005) and in combination with PACEtomo (Eisenstein *et al*, 2023). The nominal magnification was 42,000 x resulting in a pixel size of 1.069 Å in super-resolution mode. Tilt-series were collected from -60° to 60° degrees with a 3° increment, and at a defocus range of -2 µm to -7 µm, following a hybrid dose scheme (Sanchez *et al*., 2020). Hybrid tomograms had a zero-tilt image with a total dose of ∼20 e^-^/Å^2^, with the remaining dose equally distributed over the remaining images. The total electron exposure per tilt series was 100 e^-^/Å^2^ for full-length EHD2, and 158 e^-^/Å^2^ for EHD2^ΔN^. Tomograms were processed semi-automatically with tomoBEAR (Balyschew *et al*, 2023), with the key steps including MotionCorrection (Zheng *et al*, 2017), CTF determination (Zhang, 2016), fiducial-based tilt series alignment (Coray *et al*, 2024) followed by manual refinement and reconstruction by weighted back-projection in IMOD etomo (Kremer *et al*, 1996).

### Subtomogram averaging

For both constructs, subtomograms were picked along the central axis of the lipid tubules using a filament model in Dynamo catalogue (Castao-Dez *et al*., 2017) in eight-times binned tomograms (8.552 Å/pix). A total of 14,491 and 30,449 cropped points were defined for full-length EHD2 and EHD2^ΔN^, respectively. The initial coordinates of the subtomograms were imported into SUSAN (github.com/KudryashevLab/SUSAN) and reconstructed with the angular information defined by the filament model. The initial average, which showed cylindrical density, was used as a starting reference for subtomogram alignment and classification. Two iterations with only translational searches and fixed low-pass filter, followed by 10 iterations with translational and rotational searches and adaptive low-pass filter were performed. At this stage, different strategies were implemented for each construct. In the case of full-length EHD2, aligned particles were then imported into RELION-4.0 (Kimanius *et al*, 2021) and classified into 8 classes in bin8 with global angular search and 7.5° step. Four classes (6,932 particles) representing ring-like densities of different radii were selected for further processing. Particles were pooled and averaged, followed by symmetry expansion with C8 symmetry to sample the non-aligned parts of the rings, performing “subboxing” along the ring surface. Half-set IDs of subboxed particles were kept same as their respective “full-ring” particles to ensure no spurious correlations in the FSCs. Particles were then recentered on the ring surface and subjected to auto-refinement in bin8. After this, some subboxed positions converged on the same particles, leaving 44095 particles after duplicate removal. Consecutive rounds of auto-refinement followed by duplicate removal were performed on bin2 (with and without imposing local symmetry) and unbinned particles, followed by one round of polishing and CTF refinement without high-order aberrations. Final auto-refinement of polished particles in bin1 with C2 symmetry led to a 9 Å resolution map. A final cycle of TomoFrameAlignment and CTF refinement with tighter mask resulted in an 8 Å resolution map.

A final subset of 37,169 particles before polishing was then converted into a dynamo-style table (Castao-Dez *et al*, 2012) and then projected on the high-dose non-tilted images and converted to SPA-style particles STAR file. This was done with the custom script adopted from the hybridSTA (Sanchez *et al*., 2020) method. These particles were imported into CryoSPARC (Punjani *et al*, 2017) and subjected to local refinement with non-uniform filtering, angular search constrained to 1 degree and translational search to 10 Å. Then, particles were reoriented to match the C2 symmetry axis and C2 symmetry was applied in all successive alignment rounds. The consensus map of the row of 14 EHD2 monomers was used to focus on the central six monomers of EHD2 and the particle set was expanded by subboxing four monomers on each side, giving a final set of 75,439 particles after duplicate removal. Several rounds of particle subtraction, mask optimization and local refinement were performed. Lastly, the final particle stack was re-imported back into RELION (Scheres, 2012) for reconstruction, postprocessing and local resolution estimation. The final map had a nominal resolution of 6.7 Å.

In the case of EHD2^ΔN^, particle coordinates were projected on high-dose micrographs at 0° tilt using a modified hybridSTA script. Then, particles were extracted, binned 4 times and subjected to 2D classification in RELION (Scheres, 2012) into 50 classes using the default global in-plane angular search. Six classes (1,838 particles) were imported into RELION-4.0 (Kimanius *et al*., 2021), auto-refined in bin8 using a spherical mask and global angular search. Afterwards, particles were classified in 3D into 4 classes with a constrained alignment between -45° to 45°. The class (506 particles) which showed a full circle on a Z-slice was selected and auto-refined. The resulting map was used to subbox particle rows along the pseudo-helix in ChimeraX (UCSF) (Meng *et al*, 2023) using a custom script to keep particle poses. After subboxing, 17,204 particles were reoriented to match the symmetry axis, auto-refined in bin4, bin2 and unbinned with applied C2 symmetry, local angular searches and a mask covering the 12 central monomers in the row. Final auto-refinement of unbinned polished particles with a mask covering the 8 central monomers yielded a map at 13 Å resolution. These particles were then converted into a Dynamo-style table, the coordinates were projected on high-dose non-tilted images, and converted to a SPA-style particles STAR file. This was done with a custom script adopted from the hybridSTA method. These particles were imported into CryoSPARC and subjected to local refinement with non-uniform filtering, C2 symmetry, angular search constrained to 1° and translational search to 8 Å. The final map at 10.1 Å resolution includes eight central monomers. Local resolution was estimated in CryoSPARC (Punjani *et al*., 2017).

### Atom model refinement into maps

The atomic models consistent with the cryo-EM maps were generated using MDfit (Whitford *et al*, 2011). MDfit uses the cryo-EM map as an umbrella potential to bias (i.e. deform) an underlying structure-based model (SBM) (de Oliveira *et al*, 2022) in order to maximize the cross-correlation between the experimental density and the simulated electron density. An SBM is a molecular force field that is explicitly, albeit not rigidly, biased toward a certain native structure. The SBM for fitting was the EHD2 homo-dimeric crystal structure (4CID) with the sequence homology modeled by Swiss-Model (Guex & Peitsch, 1997) to remove any missing residues. The portion of the SBM for the KPF loop (residue 110-135), which is missing from the EHD2 structure, is based on the EHD4 crystal structure (pdb 5MVF). Building the SBM from the two crystal structures ensured that the resulting model was maximally consistent with the crystal conformation. This entailed no significant changes in structure as the sequences are highly similar. A preprocessing step was then necessary to move the EH domains within the dimer into a cis positioning because pdb 4CID placed the EH domains in trans. This involved only reorientation of the (421-439 loop), no other residue positions were changed. We refer to this dimeric structure as EHD2-init. An SBM using EHD2-init as the input structure was then generated using SMOGv2.3 (Noel *et al*, 2016) with the template “SBM_AA” meaning all non-hydrogen atoms were explicitly represented.

The density corresponding to the central two dimers within the cryo-EM map was chosen as the constraint for MDfit, since this region had the best resolution. Relaxation of the SBM under the influence of the cryo-EM map is performed by molecular dynamics (MD), and, thus, requires an initial condition. Two EHD2-init were rigid-body fit into the map using the “Fit in Map” tool of Chimera. This tetramer includes all studied interfaces and is in principle sufficient to model, however, the unfilled electron density due to missing filament neighbors would disrupt the fit. In order to initialize the neighbors on either side of two dimers, the translational symmetry of the filament was exploited. Four additional copies of EHD2-init were added, two positioned on either side, placed such that each dimer-dimer interface was identical. Technically, this was performed by 1) measuring the transformation X between the two central dimers in VMD, 2) duplicating the central dimers, and 3) applying X or -X to the duplicates. This six-dimer system served as the initial condition for MD. Alternating every 10******4 MD steps, 1) the dynamics were subject to only the SBM and electron density umbrella, 2) additionally a symmetrizing restraint potential. The symmetrizing restraint potential was implemented by root mean square deviation fitting a central monomer to each monomer and employing weak position restraints. This process allowed the structure to explore the cryo-EM density while additionally maintaining the symmetry of the filament. Through this iterative process, the structure converged within 3×10******5 steps. The middle two dimers were taken as the atomic model. Note that even though the filament’s local C2 rotational symmetry was not explicitly enforced by us during MD, the fact that the SBM was based on a C2 symmetric structure ensured that this symmetry was included. A final energy minimization step was performed in Phenix (Liebschner *et al*, 2019).

### Resin embedding and sectioning

Resin blocks of Human Umbilical Vein Endothelial Cells (HUVEC) were used from a previous publication (Matthaeus *et al*., 2019). In short, HUVECs were fixed using 2.5% glutaraldehyde (Sigma-Aldrich G5882-10ml) and 1% tannic acid in 0.1 M phosphate buffer pH 7.2 at room temperature for 1 hour. After fixation the cells were washed with 0.1 M phosphate buffer, scraped from the cell culture plates and pelleted by centrifugation at 1,000 x g for 5 minutes. HUVECs in suspension were embedded in 1.5% low melting agarose in Milli-Q water. The agarose block was cut into smaller cubes and processed for transmission electron microscopy. The agarose-HUVECs cubes were fixed overnight at 4 °C in 2.5% glutaraldehyde in 0.1M sodium cacodylate buffer pH 7. After washing, osmification for 2 hours was carried out at room temperature using 1% OsO4 in 0.1 M sodium cacodylate pH 7. Excess osmium was washed using MilliQ water and samples were incubated for 1 hour at 4°C in 2% uranyl acetate in MilliQ water. Dehydration was carried out using an increasing ethanol series 30% for 15 minutes, 50% for 30 minutes, 70% for 30 minutes, 90% for 30 minutes, and twice in 100% ethanol for 30 minutes each step. After dehydration, the cubes were incubated for 15 minutes in propylene oxide. Infiltration with epoxy resin (Polybed812, Polysciences) was carried out at room temperature by incubating the samples for 40 minutes in 50% and for 40 minutes in 70% resin in propylene oxide. Infiltration with 100% resin was carried out overnight at room temperature. Polymerization was carried out in for 48 hours at 60°C. Resin blocks were sectioned using a Reichert Ultracut S ultramicrotome and an Ultra 35° diamond knife (Diatome). For ultrastructural morphology assessment 70 nm sections were collected on in-house prepared Formvar/carbon 100 hexagonal mesh copper grids. Prior to collecting the 150 nm and 170 nm thick sections for electron tomography, fiducial gold beads were adsorbed onto the grid surface.

### Room-temperature tomography

Tilt series were acquired using a FEI Talos L120C electron microscope equipped with a Ceta detector and operated at 120 kV. All tilt series were collected using the software SerialEM (Mastronarde, 2005). The nominal magnification was 42,000 x resulting in a pixel size of 3.171 Å. Images were collected Images were collected from 60° to -60° with a 2° increment and bidirectionally, starting at 0°. Tomograms were reconstructed manually using IMOD (Kremer *et al*., 1996). For caveolae morphology analysis and visualization, tomograms were binned 4 times and reconstructed using a SIRT-like filter with 8 iterations. Measurements were done in IMOD. Structures of interest were segmented using Microscopy Image Browser (Belevich *et al*, 2016) and Amira (ThermoFisher Scientific). Smoothening of segmented surfaces, visualization and videos were created using Chimera (UCSF) and ChimeraX (UCSF) (Meng *et al*., 2023).

For statistics and plotting, GraphPad Prism v.7.05 was used. Normal distribution was assessed by applying a D’Agostino-Pearson test. To calculate the significant difference between two groups, normally distributed data was analyzed using a Student t test (two-tailed P-value), otherwise a Mann-Whitney-Rank-Sum (two-tailed P-value) was used. Differences of p ≤ 0.05 were considered significant (p ≤ 0.05*, p ≤ 0.01**, p ≤ 0.001***, p ≤ 0.0001****).

## Data availability

The STA-derived cryo-EM densities of the EHD2 and EHD2^ΔN^ oligomers have been deposited in the Electron Microscopy Data Bank (EMDB) under accession codes EMD-53909 (https://www.ebi.ac.uk/emdb/EMD-53909) and EMD-53911 (https://www.ebi.ac.uk/emdb/EMD-53911), respectively. Coordinates for the oligomeric EHD2 and EHD2^ΔN^ models were submitted to the Protein Data Bank (PDB) under accession codes 9RBU (https://doi.org/10.2210/pdb9RBU/pdb) and 9RCI (https://doi.org/10.2210/pdb9RCI/pdb), respectively.

## Author contributions

Elena Vázquez-Sarandeses: Conceptualization; Formal analysis; Investigation; Visualization; Writing—original draft; Writing—review and editing; EVS prepared the samples, designed and performed the experiments, analyzed the data, processed cryo-ET data, and contributed to the cryo-ET image analysis. Vasilii Mikirtumov: Formal analysis; Investigation; Visualization; Writing—review and editing. VK processed cryo-ET data and performed image analysis. Jeffrey K. Noel: Formal analysis; Investigation; Visualization; Writing—review and editing; JKN performed the flexible fitting of EHD2 into the cryo-ET density and analyzed the EH-domain fitting. Mikhail Kudryashev: Formal analysis; Supervision; Funding acquisition. Oliver Daumke: Conceptualization; Formal analysis; Supervision; Funding acquisition; Writing—original draft; Project administration; Writing—review and editing.

## Acknowledgments

We thank Dr. Thiemo Sprink, Metaxia Stavroulaki, and Dr. Christoph Diebolder from the core facility for cryo-EM at Charité Universitätsmedizin Berlin for help with the cryo-EM grid preparation and data collection. We thank Drs. Claudia Matthäus und Mara-Camelia Rusu for providing resin-embedded HUVECs. The authors thank the high-performance computing team of the MDC and Max Cluster for computational resources and Arthur Melo for constant discussions on the project.

---

**Supplementary Figure 1:**
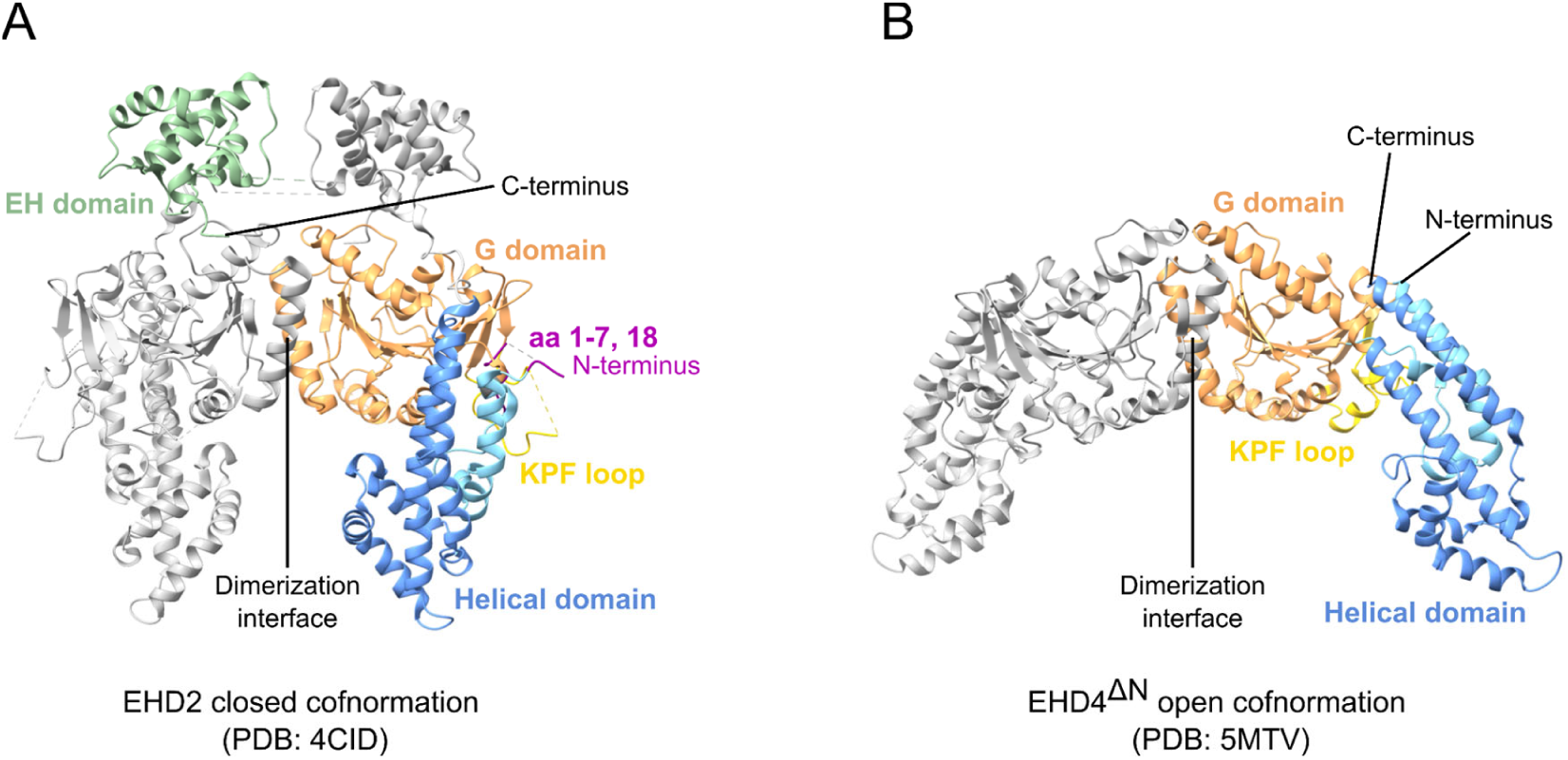
Structural overview for the open and closed EHD dimers. **A)** Crystal structure of the EHD2 dimer in the closed conformation (PDB: 4CID). One monomer is colored in gray and the other monomer is colored according to the domain architecture shown in Fig. 1A. **B)** Crystal structure of the EHD4^ΔN^ dimer in the open conformation, featuring a 50° rotation of the helical domains (PDB: 5MTV).

**Supplementary Figure 2:**
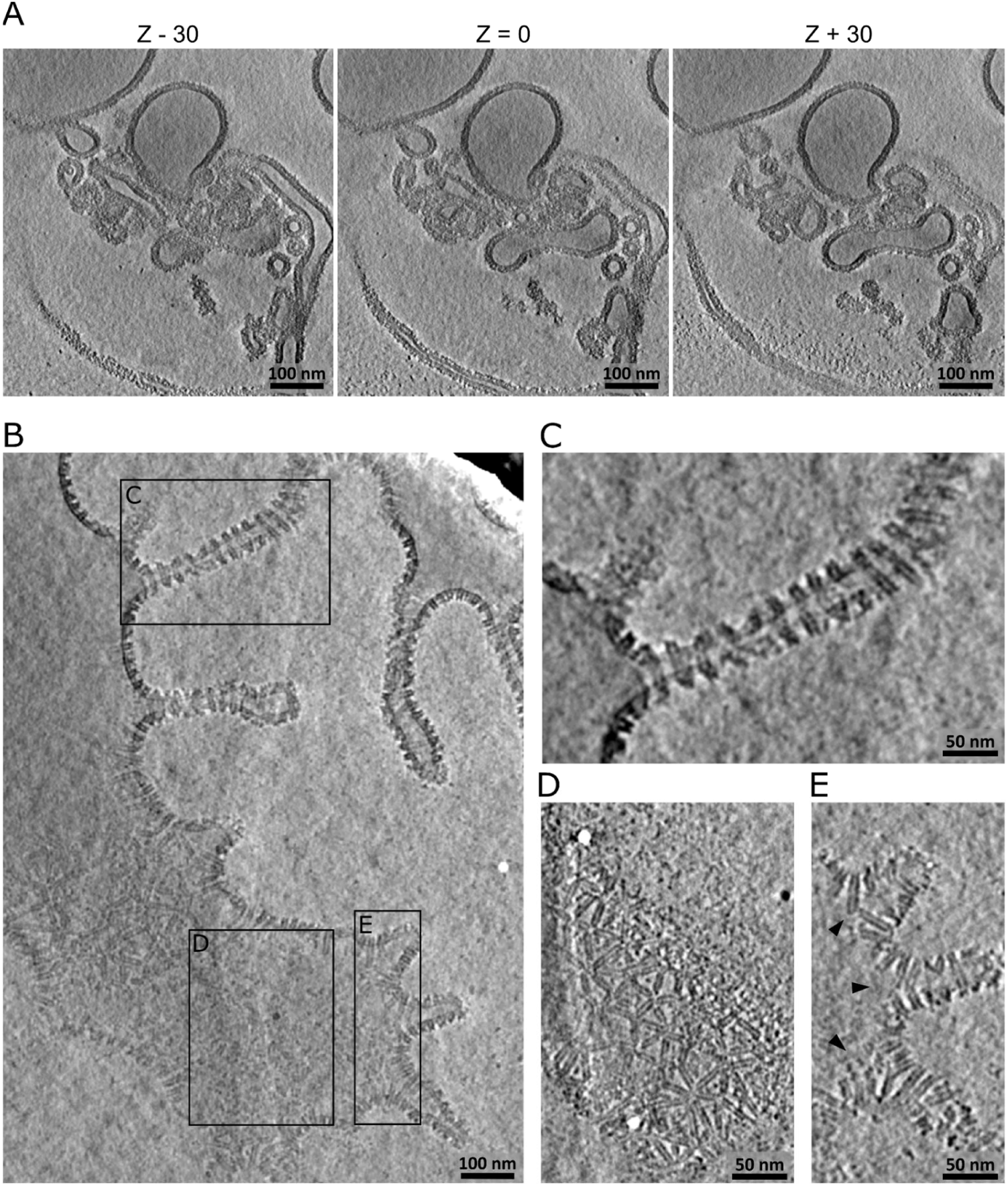
EHD2 oligomerizes on lipid bilayers of different curvature in an ATP-binding-dependent manner. **A)** Representative tomogram of nucleotide-free EHD2 reconstituted in liposomes. The middle panel shows the central Z-slice (Z = 0) in which the lumen of the lipid tubule can be observed. To show that in the apo state EHD2 cannot organize into regular filaments, other Z slices (−30 and +30) showing the surface of the lipid tubules and non-tubulated liposomes are displayed on the left and right panels, respectively. **B)** Central Z-slice of a representative tomogram showing ATP-bound EHD2 oligomeric filaments of varying lengths on the surface of lipid bilayers of different curvature. **C)** ATP-bound EHD2 ring-like oligomers on lipid tubules of high curvature. **D)** ATP-bound EHD2 oligomers on membranes of low curvature. **E)** Membrane tubulation of EHD2 in presence of ATP occurs at areas of higher curvature where oligomeric filaments encounter each other (arrowheads). Images in panels **C**, **D** and **E** correspond to the inlets highlighted in **B** and show selected Z slices of the tomographic volume.

**Supplementary Figure 3:**
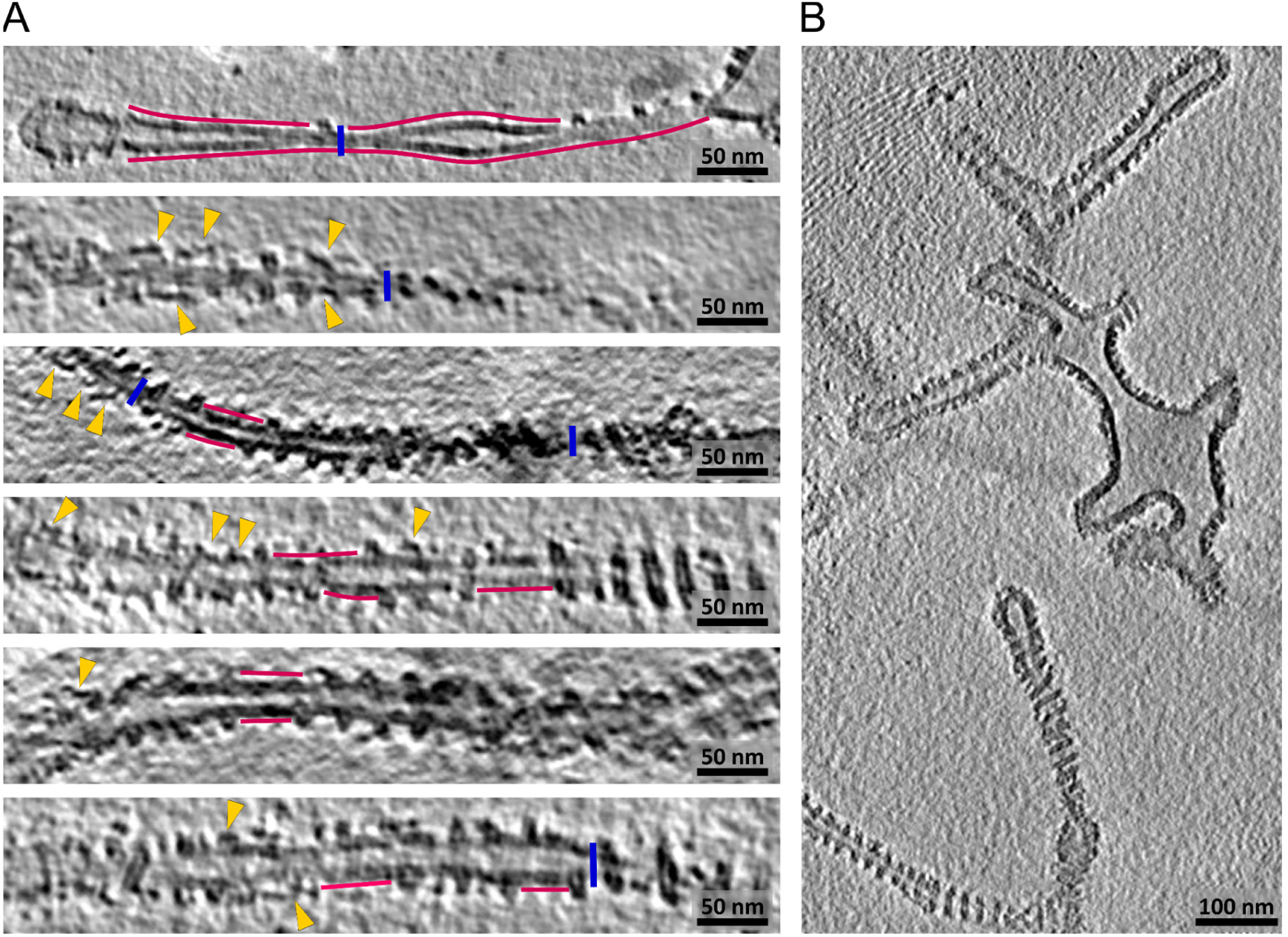
ATP hydrolysis leads to oligomer disassembly and membrane destabilization. **A)** Gallery of representative lipid tubules found in tomograms after incubating full-length EHD2 with ATP and liposomes for 120 min. At this time point, about 90% of the ATP is converted to ADP. The central slice of the tomograms is shown. Increased spacing between EHD2 oligomers, interruptions in the protein decoration or almost complete absence of protein were observed (magenta lines). Some areas of the tubules were much thinner or seemed to have collapsed (blue lines). Detached semi-open or open particles were found (yellow arrowheads). **B)** Lipid tubules with a normal EHD2 decoration were only rarely found under these conditions.

**Supplementary Figure 4:**
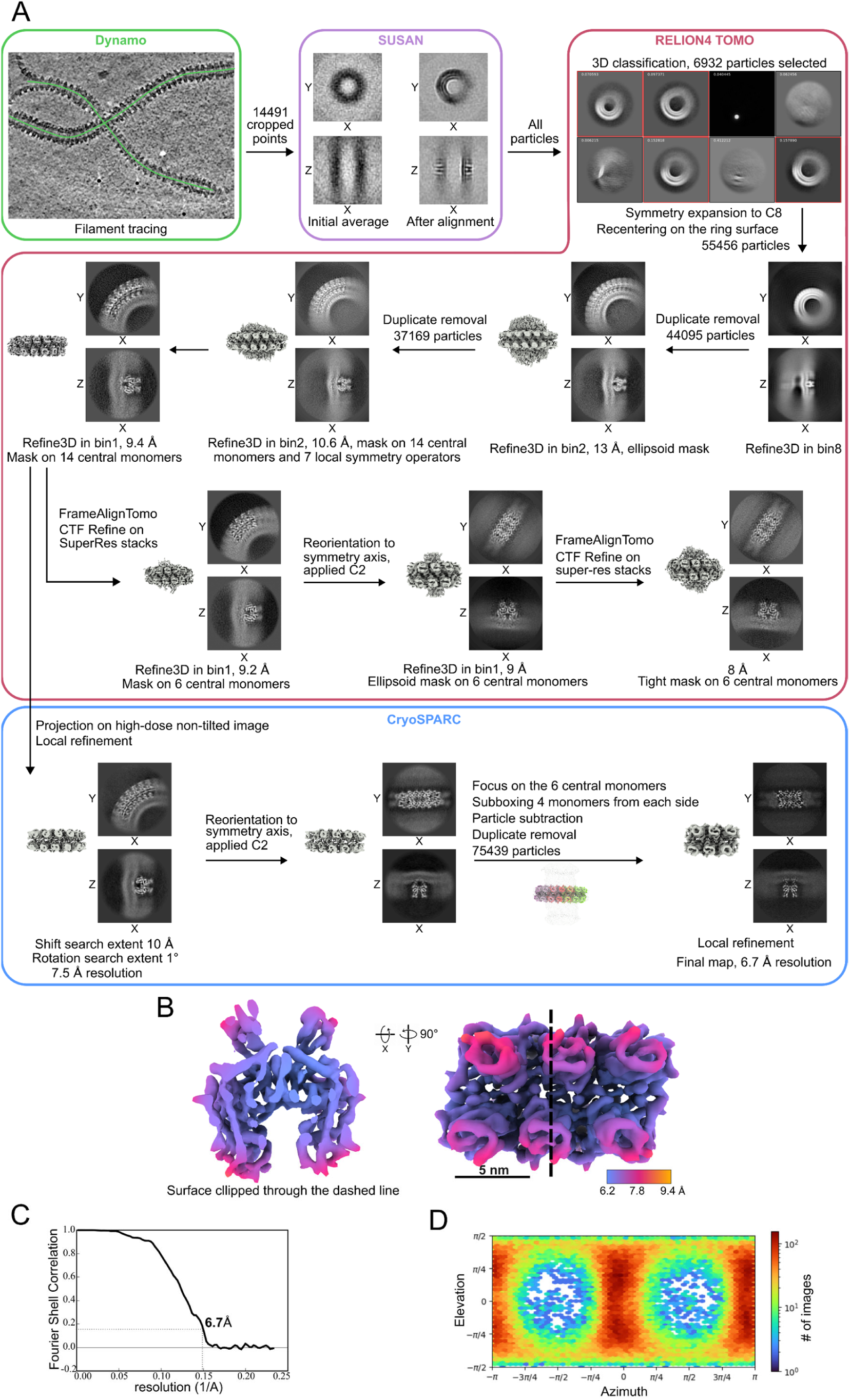
Subtomogram averaging workflow and structure determination of full-length membrane-bound EHD2. **A)** Processing flowchart, indicating prominent steps and particle number. **B)** Surface rendering of the final subtomogram averaging map colored according to local resolution. Top and front views are shown. **C)** Fourier shell correlation curve. **D)** Angular distribution plot.

**Supplementary Figure 5:**
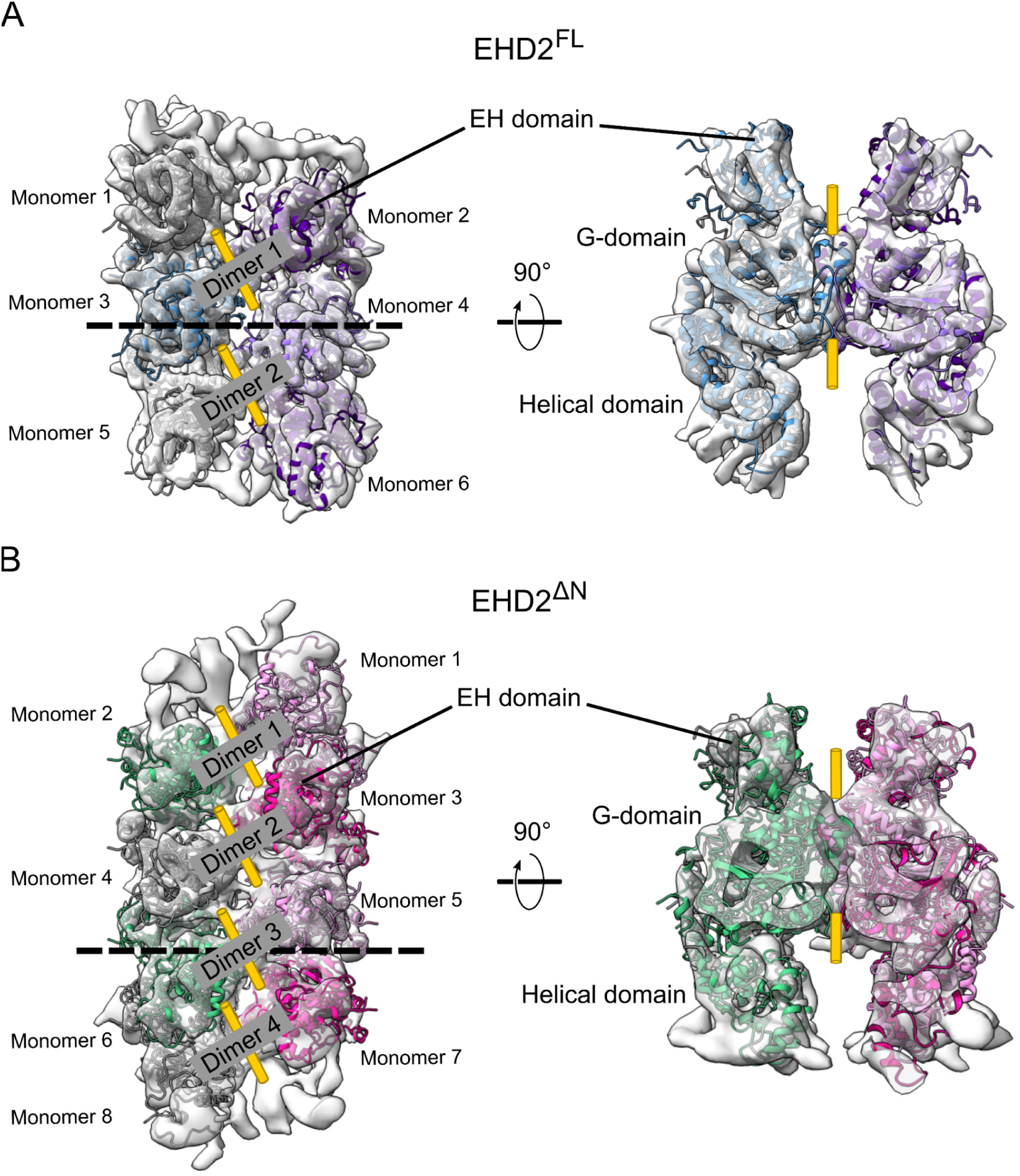
Flexible fit of the EHD2 and EHD2^ΔN^ dimers in the cryo-ET density. The closed EHD2 structure (PDB: 4CID) was fitted into the subtomogram average map of membrane-bound full-length **(A)** and N-terminally truncated **(B)** EHD2. The density of the asymmetric unit is shown overlaid with the resulting model from the top (left) and front (clipped, right) views. The dashed lines indicate where the density is clipped. The fit did not require major rearrangements, except for the EH domain and the KPF loop. The yellow tubes indicate the two-fold symmetry axes.

**Supplementary Figure 6:**
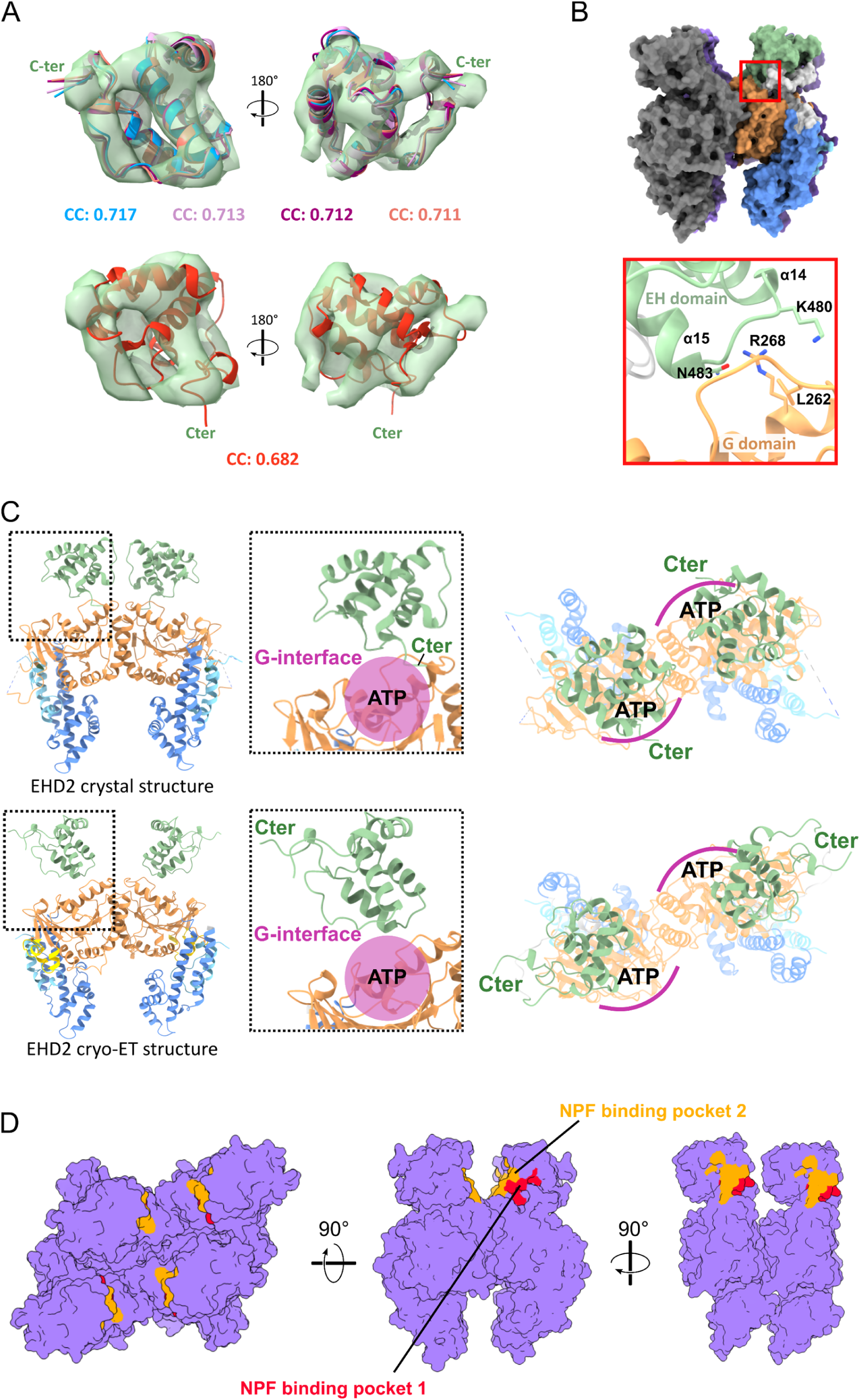
A new orientation of the EH domain. **A)** Flexible fitting of seventy different rotations of the EH domain confirms the large-scale movement. The best four scoring results, ranging correlation coefficients (CC) between the map and the model from 0.711 to 0.717, are in the rotated configuration with the C-terminal tail pointing upwards and to the outside of the filament. The crystal structure configuration with the C-terminal tail folding back to the G-domain (red cartoons) does not fit well in the cryo-ET density and resulted in a poorer CC score. Left panels: front view, same as in B. **B)** In this conformation, the EH domain might generate contacts with the G-domain directly below (red square, magnified). **C)** Top: The C-terminal tail of the EH domains folds back to the nucleotide pocket of the G domain in the EHD2 crystal structure (PDB: 4CID). It is likely that this orientation prevents the formation of the G-interface, likely representing an autoinhibited conformation that prevents oligomerization. Bottom: In the new orientation of the EH domain in the cryo-ET structure, the C-terminal tail points towards the outside the filament allowing the formation of the G-interface. The linker is not shown for visualization purposes. **D)** The NPF-binding pockets of the EH domains are buried in the filament facing inwards. Accessibility for NPF-motif containing proteins might be compromised in this configuration. Another conformation of the EH domain in complex with binding partners can therefore be envisaged.

**Supplementary Figure 7:**
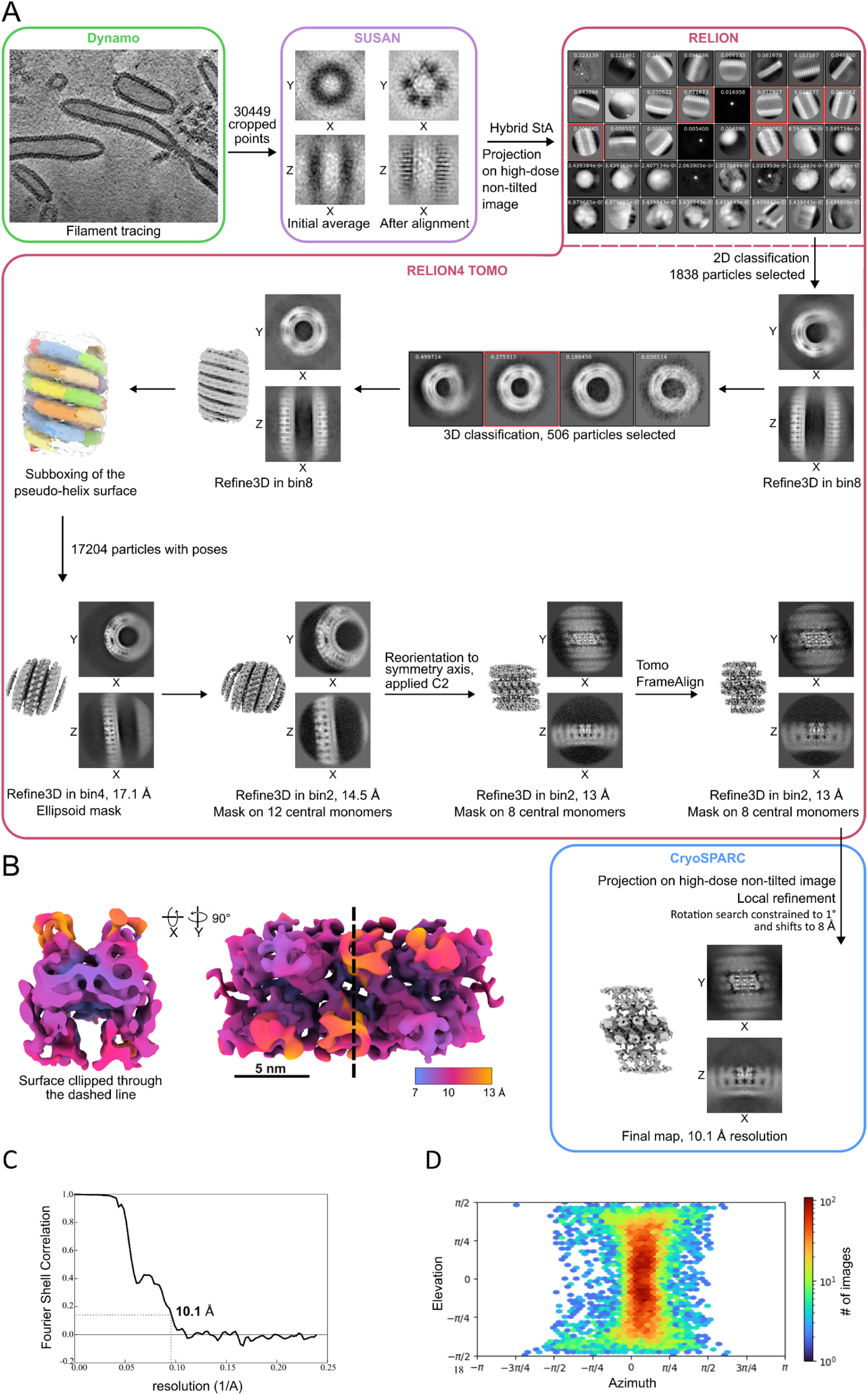
Subtomogram averaging workflow and structure determination of N-terminally truncated membrane-bound EHD2. **A)** Processing flowchart, indicating prominent steps and particle number. **B)** Surface rendering of the final subtomogram averaging map colored according to local resolution. Top and front views are shown. **C)** Fourier Shell Correlation curve. **D)** Angular distribution plot.

**Supplementary Table 1:**
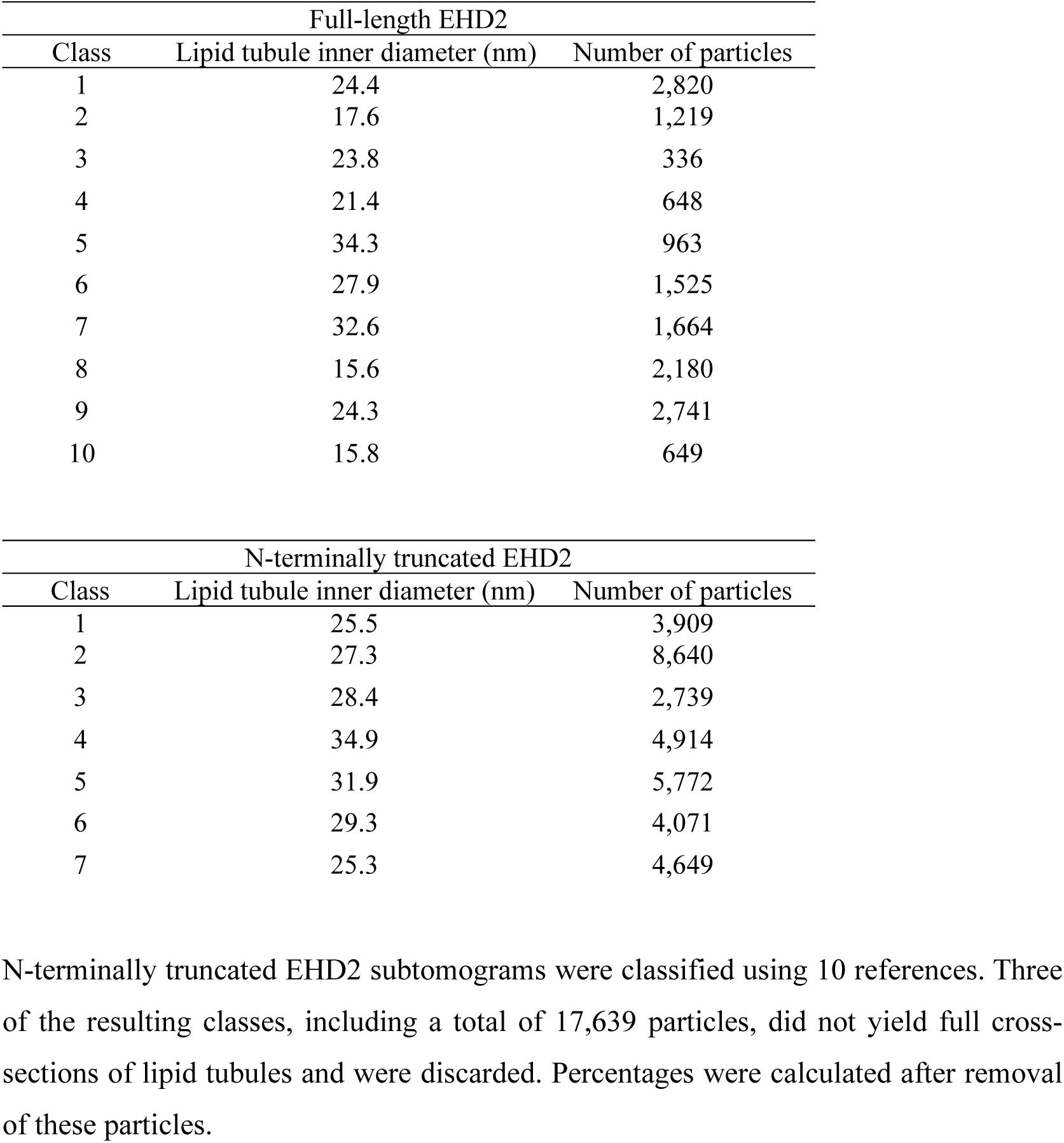
Multireference classification and distribution of lipid tubule diameters for full-length and N-terminally truncated EHD2

**Supplementary Table 2:** Data collection, refinement and validation statistics

|  | EHD2 | EHD2ΔN |
| --- | --- | --- |
| **Data collection and processing** |  |  |
| Magnification | 42,000 | 42,000 |
| Voltage (kV) | 300 | 300 |
| Electron exposure (e<sup>-</sup>/Å<sup>2</sup>) | 100 | 158 |
| Defocus range (μm) | -2 – -7 | -1.5 – -5 |
| Pixel size (Å) | 1.069 | 1.069 |
| Symmetry imposed | C2 | C2 |
| Initial particle images (no.) | 14,491 | 30,449 |
| Final particle images (no.) | 6,932 | 17,204 |
| Map resolution (Å) | 6.7 | 11 |
| FSC threshold | 0.143 | 0.143 |
| Map resolution range (Å) | 6.2 – 9.4 | 6.3 – 16.6 |
| **Refinement** |  |  |
| Initial model used (PDB code) | 4CID | 4CID |
| Map sharpening *B* factor (Å<sup>2</sup>) | -200 | -200 |
| Model composition |  |  |
| Non-hydrogen atoms | 16,576 | 16,576 |
| Protein residues | 2084 | 2084 |
| Ligands | 0 | 0 |
| R.m.s. deviations |  |  |
| Bond lengths (Å) | 0.003 | 0.003 |
| Bond angles (°) | 0.880 | 0.895 |
| Validation |  |  |
| MolProbity score<sup>1</sup> | 1.31 | 1.29 |
| Clashcore | 2.90 | 3.57 |
| Poor rotamers (%) | 0.28 | 0.23 |
| Ramachandran plot |  |  |
| Favored (%) | 96.6 | 97.3 |
| Allowed (%) | 3.2 | 2.4 |
| Disallowed (%) | 0.2 | 0.4 |

<sup>1</sup> (Williams *et al*, 2018)

**Supplementary Table 3:**
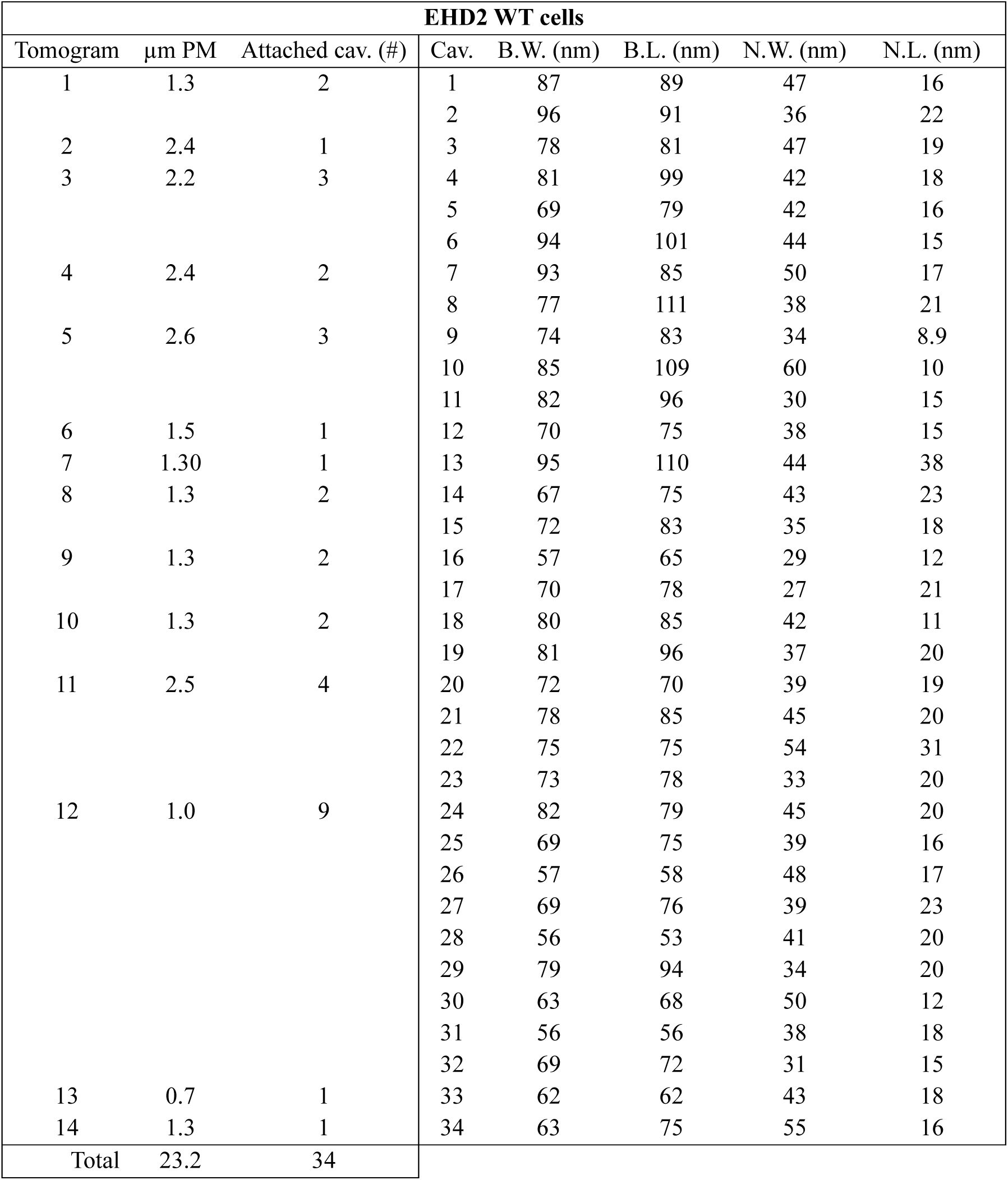
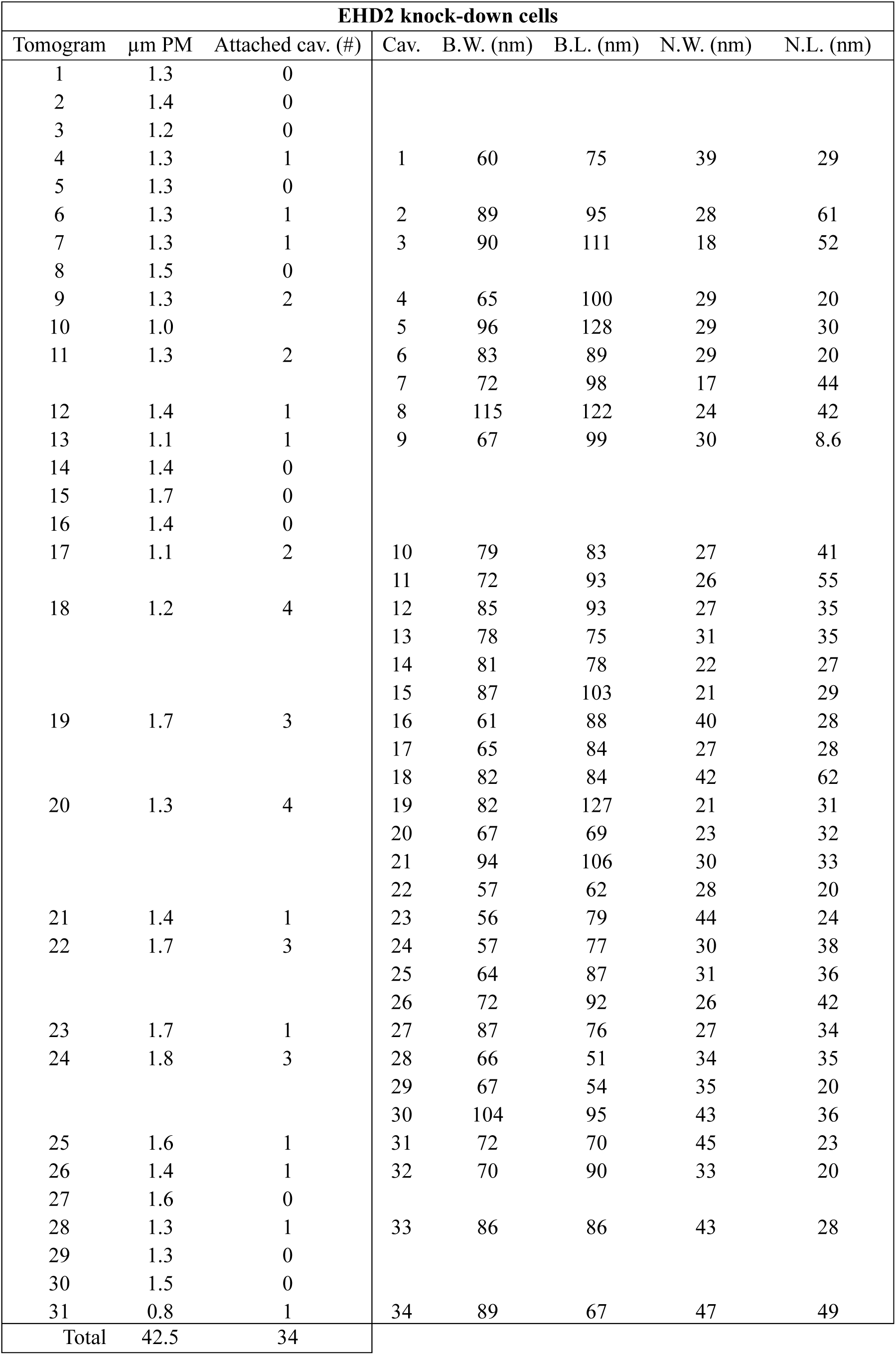
Analysis of caveolae morphology in the presence and absence of EHD2. Only caveolae (cav.) still connected to the plasma membrane (PM), e.g. attached caveolae were considered. The bulb width (B.W.), bulb length (B.L.), neck width (N.W.) and neck length (N.L.) were measured.

## References

1. Balyschew N, Yushkevich A, Mikirtumov V, Sanchez RM, Sprink T, Kudryashev M (2023) Streamlined structure determination by cryo-electron tomography and subtomogram averaging using TomoBEAR. Nat Comm 14: 6543, doi: 10.1038/s41467-023-42085-w

2. Belevich I, Joensuu M, Kumar D, Vihinen H, Jokitalo E (2016) Microscopy Image Browser: A Platform for Segmentation and Analysis of Multidimensional Datasets. PLoS Biol 14: e1002340, doi: 10.1371/journal.pbio.1002340

3. Bhattacharyya S, Pucadyil TJ (2020) Cellular functions and intrinsic attributes of the ATP-binding Eps15 homology domain-containing proteins. Prot Sci 29: 1321–1330, doi: 10.1002/pro.3860

4. Campelo F, Fabrikant G, McMahon HT, Kozlov MM (2010) Modeling membrane shaping by proteins: Focus on EHD2 and N-BAR domains. FEBS Letters 584: 1830–1839 doi: 10.1016/j.febslet.2009.10.023

5. Caplan S, Naslavsky N, Hartnell LM, Lodge R, Polishchuk RS, Donaldson JG, Bonifacino JS (2002) A tubular EHD1-containing compartment involved in the recycling of major histocompatibility complex class I molecules to the plasma membrane. EMBO J 21: 2557–2567, doi: 10.1093/emboj/21.11.2557

6. Castao-Dez D, Kudryashev M, Arheit M, Stahlberg H (2012) Dynamo: A flexible, user-friendly development tool for subtomogram averaging of cryo-EM data in high-performance computing environments. J Struct Biol 178: 139–151, doi: 10.1016/j.jsb.2011.12.017

7. Castao-Dez D, Kudryashev M, Stahlberg H (2017) Dynamo Catalogue: Geometrical tools and data management for particle picking in subtomogram averaging of cryo-electron tomograms. J Struct Biol 197: 135–144, doi: 10.1016/j.jsb.2016.06.005

8. Coray R, Navarro P, Scaramuzza S, Stahlberg H, Castaño-Díez D (2024) Automated fiducial-based alignment of cryo-electron tomography tilt series in Dynamo. Structure 32: 1808–1819, doi: 10.1016/j.str.2024.07.003

9. Curran J, Makara MA, Little SC, Musa H, Liu B, Wu X, Polina I, Alecusan JS, Wright P, Li J et al (2014) EHD3-dependent endosome pathway regulates cardiac membrane excitability and physiology. Circ Res 115: 68–78 doi: 10.1161/CIRCRESAHA.115.304149

10. Daumke O, Lundmark R, Vallis Y, Martens S, Butler PJG, McMahon HT (2007) Architectural and mechanistic insights into an EHD ATPase involved in membrane remodelling. Nature 449: 923–927, doi: 10.1038/nature06173

11. Daumke O, Praefcke GJ (2016) Invited review: Mechanisms of GTP hydrolysis and conformational transitions in the dynamin superfamily. Biopolymers 105: 580–593, doi: 10.1002/bip.22855

12. de Oliveira AB, Jr., Contessoto VG, Hassan A, Byju S, Wang A, Wang Y, Dodero-Rojas E, Mohanty U, Noel JK, Onuchic JN et al (2022) SMOG 2 and OpenSMOG: Extending the limits of structure-based models. Prot Sci 31: 158–172, doi: 10.1002/pro.4209

13. Demonbreun AR, Swanson KE, Rossi AE, Deveaux HK, Earley JU, Allen MV, Arya P, Bhattacharyya S, Band H, Pytel P et al (2015) Eps 15 homology domain (EHD)-1 remodels transverse tubules in skeletal muscle. PLoS ONE 10: e0136679, doi: 10.1371/journal.pone.0136679

14. Deo R, Kushwah MS, Kamerkar SC, Kadam NY, Dar S, Babu K, Srivastava A, Pucadyil TJ (2018) ATP-dependent membrane remodeling links EHD1 functions to endocytic recycling. Nat Comm 9doi: 10.1038/s41467-018-07586-z

15. Doherty KR, Demonbreun AR, Wallace GQ, Cave A, Posey AD, Heretis K, Pytel P, McNally EM (2008) The endocytic recycling protein EHD2 interacts with myoferlin to regulate myoblast fusion. J Biol Chem 283: 20252–20260, doi: 10.1074/jbc.M802306200

16. Eisenstein F, Yanagisawa H, Kashihara H, Kikkawa M, Tsukita S, Danev R (2023) Parallel cryo electron tomography on in situ lamellae. Nat Methods 20: 131–138 doi: 10.1038/s41592-022-01690-1

17. Giridharan SSP, Cai B, Vitale N, Naslavsky N, Caplan S (2013) Cooperation of MICAL-L1, syndapin2, and phosphatidic acid in tubular recycling endosome biogenesis. Mol Biol Cell 24: 1776–1790, doi: 10.1091/mbc.E13-01-0026

18. Guex N, Peitsch MC (1997) SWISS-MODEL and the Swiss-PdbViewer: an environment for comparative protein modeling. Electrophoresis 18: 2714–2723, doi: 10.1002/elps.1150181505

19. Guilherme A, Soriano NA, Furcinitti PS, Czech MP (2004) Role of EHD1 and EHBP1 in perinuclear sorting and insulin-regulated GLUT4 recycling in 3T3-L1 adipocytes. J Biol Chem 279: 40062–40075, doi: 10.1074/jbc.M401918200

20. Hoernke M, Mohan J, Larsson E, Blomberg J, Kahra D, Westenhoff S, Schwieger C, Lundmark R (2017) EHD2 restrains dynamics of caveolae by an ATP-dependent, membrane-bound, open conformation. Proc Natl Acad Sci U S A: E4360–E4369, doi: 10.1073/pnas.1614066114

21. Hubert M, Larsson E, Vegesna NVG, Ahnlund M, Johansson AI, Moodie LWK, Lundmark R (2020) Lipid accumulation controls the balance between surface connection and scission of caveolae. eLife 9: 1–31, doi: 10.7554/eLife.55038

22. Jones T, Naslavsky N, Caplan S (2020) Eps15 Homology Domain Protein 4 (EHD4) is required for Eps15 Homology Domain Protein 1 (EHD1)-mediated endosomal recruitment and fission. PLoS ONE 15: e0239657, doi: 10.1371/journal.pone.0239657

23. Kieken F, Jović M, Tonelli M, Naslavsky N, Caplan S, Sorgen PL (2009) Structural insight into the interaction of proteins containing NPF, DPF, and GPF motifs with the C-terminal EH-domain of EHD1. Protein Sci 18: 2471–2479, doi: 10.1002/pro.258

24. Kimanius D, Dong L, Sharov G, Nakane T, Scheres SHW (2021) New tools for automated cryo-EM single-particle analysis in RELION-4.0. Biochem J 478: 4169–4185 doi: 10.1042/BCJ20210708

25. Kong L, Sochacki KA, Wang H, Fang S, Canagarajah B, Kehr AD, Rice WJ, Strub MP, Taraska JW, Hinshaw JE (2018) Cryo-EM of the dynamin polymer assembled on lipid membrane. Nature 560: 258–262, doi: 10.1038/s41586-018-0378-6

26. Kozlov MM, Taraska JW (2023) Generation of nanoscopic membrane curvature for membrane trafficking. Nat Rev Mol Cell Biol 24: 63–78 doi: 10.1038/s41580-022-00511-9

27. Kremer JR, Mastronarde DN, McIntosh JR (1996) Computer Visualization of Three-Dimensional Image Data Using IMOD. J Struc Biol 116: 71–76, doi: 10.1006/jsbi.1996.0013

28. Lee DW, Zhao X, Scarselletta S, Schweinsberg PJ, Eisenberg E, Grant BD, Greene LE (2005) ATP binding regulates oligomerization and endosome association of RME-1 family proteins. J Biol Chem 280: 17213–17220, doi: 10.1074/jbc.M412751200

29. Liebschner D, Afonine PV, Baker ML, Bunkóczi G, Chen VB, Croll TI, Hintze B, Hung LW, Jain S, McCoy AJ et al (2019) Macromolecular structure determination using X-rays, neutrons and electrons: recent developments in Phenix. Acta Crystallogr D Struct Biol 75: 861–877, doi: 10.1107/s2059798319011471

30. Lu Q, Insinna C, Ott C, Stauffer J, Pintado PA, Rahajeng J, Baxa U, Walia V, Cuenca A, Hwang Y-S et al (2015) Early steps in primary cilium assembly require EHD1/EHD3-dependent ciliary vesicle formation. Nat Cell Biol 17: 228–240, doi: 10.1038/ncb3109

31. Ludwig A, Howard G, Mendoza-Topaz C, Deerinck T, Mackey M, Sandin S, Ellisman MH, Nichols BJ (2013) Molecular Composition and Ultrastructure of the Caveolar Coat Complex. PLoS Biol 11: e1001640, doi: 10.1371/journal.pbio.1001640

32. Malinova TS, Angulo-Urarte A, Nchel J, Tauber M, van der Stoel MM, Janssen V, de Haan A, Groenen AG, Tebbens M, Graupera M et al (2021) A junctional PACSIN2/EHD4/MICAL-L1 complex coordinates VE-cadherin trafficking for endothelial migration and angiogenesis. Nat Comm 12: 2610, doi: 10.1038/s41467-021-22873-y

33. Marg A, Schoewel V, Timmel T, Schulze A, Shah C, Daumke O, Spuler S (2012) Sarcolemmal Repair Is a Slow Process and Includes EHD2. Traffic 13: 1286–1294, doi: 10.1111/j.1600-0854.2012.01386.x

34. Mastronarde DN (2005) Automated electron microscope tomography using robust prediction of specimen movements. J Struc Biol 152: 36–51, doi: 10.1016/j.jsb.2005.07.007

35. Matthaeus C, Lahmann I, Kunz S, Jonas W, Melo AA, Lehmann M, Larsson E, Lundmark R, Kern M, Blher M et al (2020) EHD2-mediated restriction of caveolar dynamics regulates cellular fatty acid uptake. Proc Natl Acad Sci U S A 117: 7471–7481, doi: 10.1073/pnas.1918415117

36. Matthaeus C, Lian X, Kunz S, Lehmann M, Zhong C, Bernert C, Lahmann I, Mller DN, Gollasch M, Daumke O (2019) ENOS-NO-induced small blood vessel relaxation requires EHD2-dependent caveolae stabilization. PLoS ONE 14: 1–22, doi: 10.1371/journal.pone.0223620

37. Matthaeus C, Sochacki KA, Dickey AM, Puchkov D, Haucke V, Lehmann M, Taraska JW (2022) The molecular organization of differentially curved caveolae indicates bendable structural units at the plasma membrane. Nat Comm 13: 7234, doi: 10.1038/s41467-022-34958-3

38. Melo AA, Hegde BG, Shah C, Larsson E, Isas JM, Kunz S, Lundmark R, Langen R, Daumke O (2017) Structural insights into the activation mechanism of dynamin-like EHD ATPases. Proc Natl Acad Sci U S A 2: 5629–5634, doi: 10.1073/pnas.1614075114

39. Melo AA, Sprink T, Noel JK, Vzquez-Sarandeses E, van Hoorn C, Mohd S, Loerke J, Spahn CMT, Daumke O (2022) Cryo-electron tomography reveals structural insights into the membrane remodeling mode of dynamin-like EHD filaments. Nat Comm 13: 7641, doi: 10.1038/s41467-022-35164-x

40. Meng EC, Goddard TD, Pettersen EF, Couch GS, Pearson ZJ, Morris JH, Ferrin TE (2023) UCSF ChimeraX: Tools for structure building and analysis. Prot Sci 32: e4792, doi: 10.1002/pro.4792

41. Moren B, Hansson B, Negoita F, Fryklund C, Lundmark R, Gransson O, Stenkula KG (2019) EHD2 regulates adipocyte function and is enriched at cell surface-associated lipid droplets in primary human adipocytes. Mol Biol Cell 30: 1147–1159, doi: 10.1091/mbc.E18-10-0680

42. Moren B, Shah C, Howes MT, Schieber NL, McMahon HT, Parton RG, Daumke O, Lundmark R (2012) EHD2 regulates caveolar dynamics via ATP-driven targeting and oligomerization. Mol Biol Cell 23: 1316–1329 doi: 10.1091/mbc.E11-09-0787

43. Naslavsky N, Caplan S (2011) EHD proteins : key conductors of endocytic transport. Trends Cell Biol 21: 122–131, doi: 10.1016/j.tcb.2010.10.003

44. Noel JK, Levi M, Raghunathan M, Lammert H, Hayes RL, Onuchic JN, Whitford PC (2016) SMOG 2: A Versatile Software Package for Generating Structure-Based Models. PLoS Comp Biol 12: e1004794, doi: 10.1371/journal.pcbi.1004794

45. Nyenhuis SB, Wu X, Strub M-P, Yim Y-I, Stanton AE, Baena V, Syed ZA, Canagarajah B, Hammer JA, Hinshaw JE (2023) OPA1 helical structures give perspective to mitochondrial dysfunction. Nature 620: 1109–1116 doi: 10.1038/s41586-023-06462-1

46. Pant S, Sharma M, Patel K, Caplan S, Carr CM, Grant BD (2009) AMPH-1/Amphiphysin/Bin1 functions with RME-1/Ehd1 in endocytic recycling. Nat Cell Biol 11: 1399–1410, doi: 10.1038/ncb1986

47. Parton RG, Kozlov MM, Ariotti N (2020) Caveolae and lipid sorting: Shaping the cellular response to stress. J Cell Biol 219: e201905071, doi: 10.1083/JCB.201905071

48. Peng R, Rochon K, Hutson AN, Stagg SM, Mears JA (2025) The structure of the Drp1 lattice on membrane. J Mol Biol 437: 169125, doi: 10.1016/j.jmb.2025.169125

49. Posey AD, Jr., Swanson KE, Alvarez MG, Krishnan S, Earley JU, Band H, Pytel P, McNally EM, Demonbreun AR (2014) EHD1 mediates vesicle trafficking required for normal muscle growth and transverse tubule development. Dev Biol 387: 179–190, doi: 10.1016/j.ydbio.2014.01.004

50. Punjani A, Rubinstein JL, Fleet DJ, Brubaker MA (2017) CryoSPARC: Algorithms for rapid unsupervised cryo-EM structure determination. Nat Methods 14: 290–296, doi: 10.1038/nmeth.4169

51. Sanchez RM, Zhang Y, Chen W, Dietrich L, Kudryashev M (2020) Subnanometer-resolution structure determination in situ by hybrid subtomogram averaging - single particle cryo-EM. Nat Comm 11: 3709, doi: 10.1038/s41467-020-17466-0

52. Scheres SHW (2012) RELION: Implementation of a Bayesian approach to cryo-EM structure determination. J Struct Biol 180: 519–530, doi: 10.1016/j.jsb.2012.09.006

53. Senju Y, Itoh Y, Takano K, Hamada S, Suetsugu S (2011) Essential role of PACSIN2/syndapin-II in caveolae membrane sculpting. J Cell Sci 124: 2032–2040, doi: 10.1242/jcs.086264

54. Shah C, Hegde BG, Morn B, Behrmann E, Mielke T, Moenke G, Spahn CMT, Lundmark R, Daumke O (2014) Structural insights into membrane interaction and caveolar targeting of dynamin-like EHD2. Structure 22: 409–420, doi: 10.1016/j.str.2013.12.015

55. Shao Y, Akmentin W, Toledo-Aral JJ, Rosenbaum J, Valdez G, Cabot JB, Hilbush BS, Halegoua S (2002) Pincher, a pinocytic chaperone for nerve growth factor/TrkA signaling endosomes. J Cell Biol 157: 679–691, doi: 10.1083/jcb.200201063

56. Sharma M, Giridharan SSP, Rahajeng J, Caplan S, Naslavsky N (2010) MICAL-L1: An unusual Rab effector that links EHD1 to tubular recycling endosomes. Commun Integr Biol 3: 181–183, doi: 10.1091/mbc.E09-06-0535

57. Solinger JA, Rashid H-O, Prescianotto-Baschong C, Spang A (2020) FERARI is required for Rab11-dependent endocytic recycling. Nat Cell Biol 22: 213–224, doi: 10.1038/s41556-019-0456-5

58. Sotodosos-Alonso L, Pulgar M, Pozo MA (2023) Caveolae Mechanotransduction at the Interface between Cytoskeleton. Cells 12: 942, doi: 10.3390/cells12060942

59. Stoeber M, Stoeck IK, Hänni C, Bleck CK, Balistreri G, Helenius A (2012) Oligomers of the ATPase EHD2 confine caveolae to the plasma membrane through association with actin. EMBO J 31: 2350–2364, doi: 10.1038/emboj.2012.98

60. Valdez G, Akmentin W, Philippidou P, Kuruvilla R, Ginty DD, Halegoua S (2005) Pincher-mediated macroendocytosis underlies retrograde signaling by neurotrophin receptors. J Neurosci 25: 5236–5247, doi: 10.1523/JNEUROSCI.5104-04.2005

61. von der Malsburg A, Sapp GM, Zuccaro KE, von Appen A, Moss FR, 3rd, Kalia R, Bennett JA, Abriata LA, Dal Peraro M, van der Laan M et al (2023) Structural mechanism of mitochondrial membrane remodelling by human OPA1. Nature 620: 1101–1108, doi: 10.1038/s41586-023-06441-6

62. Whitford PC, Ahmed A, Yu Y, Hennelly SP, Tama F, Spahn CMT, Onuchic JN, Sanbonmatsu KY (2011) Excited states of ribosome translocation revealed through integrative molecular modeling. Proc Natl Acad Sci U S A 108: 18943–18948, doi: 10.1073/pnas.1108363108

63. Williams CJ, Headd JJ, Moriarty NW, Prisant MG, Videau LL, Deis LN, Verma V, Keedy DA, Hintze BJ, Chen VB et al (2018) MolProbity: More and better reference data for improved all-atom structure validation. Protein Sci 27: 293–315, doi: 10.1002/pro.3330

64. Zhang K (2016) Gctf: Real-time CTF determination and correction. J Struct Biol 193: 1–12, doi: 10.1016/j.jsb.2015.11.003

65. Zheng SQ, Palovcak E, Armache J-P, Verba KA, Cheng Y, Agard DA (2017) MotionCor2: anisotropic correction of beam-induced motion for improved cryo-electron microscopy. Nat Methods 14: 331–332, doi: 10.1038/nmeth.4193

